# Multi-cell type deconvolution using a probabilistic model for single-molecule DNA methylation haplotypes

**DOI:** 10.1101/2023.08.20.554012

**Authors:** I. Unterman, D. Avrahami, E. Katsman, T.J. Triche, B. Glaser, B.P. Berman

## Abstract

**Background:** Deconvolution is used to estimate the proportion of mixed cell types from tissue or blood samples based on genomic profiling. DNA methylation is commonly used because specific CpG positions reflect cell type identity and can be accurately measured at either the population or single-molecule level. Methylation sequencing techniques can profile multiple individual CpGs on a single DNA molecule, but few deconvolution models have been developed to exploit these single-molecule *methylation haplotypes* for cell type deconvolution.

**Results and Conclusions:** We used simulated whole-genome methylation data and *in silico* mixtures of real data to compare existing deconvolution tools with two new models developed here. We found that adapting an existing model *CelFiE* to incorporate methylation haplotype information improved deconvolution accuracy by ∼30% over other tools, including the original CelFiE. In addition to overall higher accuracy, our new tool CelFiE Integrated Single-molecule Haplotypes (or *CelFiE-ISH*) outperformed others in detecting rare cell types present at 0.1% and below. Detection of rare cell types is important for the analysis of circulating DNA, which we demonstrate using a patient-derived plasma sequencing dataset.Finally,we show that marker selection strategy has a strong effect on deconvolution accuracy, concluding that haplotype-aware deconvolution can take advantage of markers tailored for that purpose.

## Background

DNA methylation is an essential cell type marker [1], with patterns that are formed and maintained during development and that continue to influence gene expression patterns of differentiated tissues [2]. Alteration of DNA methylation is characteristic of several human diseases, including imprinting related syndromes like Beckwith-Wiedemann and Prader-Willi. DNA methylation abnormalities are a hallmark of cancer, and may play a role in early events leading to transformation [3].

Accurate mapping of the epigenetic landscapes of healthy and diseased cells is necessary to fully characterize the role of methylation alterations in disease pathophysiology. In primary tissues, disease-specific methylation alterations are confounded by cell type composition and intra-cell-type heterogeneity [4]. In most cases, disease samples include adjacent healthy cells as well as contamination of other cell types.

Aberrant methylation can also be detected in DNA fragments released from the primary tissue into the circulation, which occurs after cell death. Clinical testing based on cell-free DNA (cfDNA) in plasma is extensively used for prenatal screening for aneuploidy [5], and increasingly in detection, classification, and monitoring of cancer [6]. Comprehensive cell-of-origin deconvolution of circulating cfDNA can be determined using DNA methylation patterns [7], and these can be used for early detection of tissue damage, diagnosis of cancer and autoimmune disorders, and monitoring of treatment response and recurrence [8–11]. Due to the growing interest in utilizing DNA methylation profiles in clinical settings, it has become increasingly important to avoid confounding factors.

Deconvolution algorithms run the gamut from fully reference-free to reference-based [12]. Reference-free algorithms either use a predetermined number of cell types or infer the number of cell types from bulk data, but cannot name the cells or distinguish different cell types from intra-cell type heterogeneity. Semi-reference free are methods which use some known cell types (like immune cells) to subtract them out. Finally, reference-based algorithms require an input atlas with methylation patterns from known healthy and diseased samples. These methods generally depend on the purity of the reference atlas samples, and are limited to analyzing samples contained within this atlas [13]. The choice of deconvolution method also depends on the type of methylation data being used. Affinity-enrichment methods capture overall methylation levels across a region, while bisulfite-conversion and enzymatic conversion methods provide information at a single-CpG resolution [14]. Third-generation long-read sequencing technologies from Oxford Nanopore and Pacific Biosciences also yield single base-pair level DNA methylation [15,16].

Because whole-genome cell-type atlases of DNA methylation have not been available until recently, prior tools have focused on either array-based analysis [7,17–19], or analysis of specific tissue types [8,20]. CelFiE [21] was developed for whole-genome bisulfite sequencing (WGBS), and improved accuracy by incorporating the absolute number of unmethylated and methylated reads at each CpG to account for sequencing depth. None of these methods took advantage of multi-CpG patterns within individual reads, or *methylation haplotypes [22]*, which have been used in the identification of imprinted regions [23] and cancer DNA [24]. Loyfer et al. recently published the first high-quality WGBS cell type atlas of human DNA methylation, and developed their own haplotype-based deconvolution model, UXM [1]. The availability of this high resolution cell-type map is a landmark resource for the field, which we use to benchmark the performance of the publicly available deconvolution models here. Additionally, we analyze the performance of individual marker regions and show that haplotype-aware deconvolution methods should use markers tailored for methylation haplotypes.

## Results

### Algorithms overview

Each of the algorithms investigated starts with a set of cell-type specific marker regions across the genome. The CelFiE algorithm [21] takes in a reference atlas with methylation probabilities, i.e. beta values, for each cell type at each CpG site within the marker regions, and models each of the *n* cell types as an independent methylation state using these beta values (Table 1, “Num methylation states”). Each CpG within the marker region is treated independently as input to an Expectation-Maximization mixture model which attempts to assign the probability of each read originating from each of the *n* atlas cell types. An optional setting in the CelFiE package (“CelFiE-sum”) allows the atlas beta values of all CpGs within a marker region to be averaged, thus enforcing a uniform methylation level within each marker (Table 1, “Uniform methylation”). CelFiE and CelFiE-sum require input samples to contain both the number of methylated reads and total read counts for every CpG in the marker region. While this does allow the Expectation-Maximization model of CelFiE to estimate sampling variance, it does not take advantage of the single-molecule nature of bisulfite sequencing, which includes linked methylation states for multiple CpGs within a read, i.e. a methylation *haplotype* for each read [22] (Table 1, “Methylation haplotypes”). CelFiE and CelFiE-sum also re-estimate cell type methylation states in the reference atlas, in a combined analysis that includes both the atlas and test samples.

**Table 1:** Deconvolution algorithms. A comparison of the algorithms investigated, based on the inclusion of methylation haplotypes, the number of methylation states, assumption of uniformity across methylation blocks and ability to re-estimate the reference methylation probabilities.

|  | CelFiE | CeFiE Sum | UXM | CelFiE-ISH | CelFiE-ISH Reatlas | Epistate |
| --- | --- | --- | --- | --- | --- | --- |
| ref | Caggiano | Caggiano | Loyfer | this study | this study | this study |
| Methylation haplotypes | No | No | Yes | Yes | Yes | Yes |
| Num methylation states | $n$ | $n$ | 2 | $n$ | $n$ | 2 |
| Uniform methylation | No | Yes | Yes | No | No | No |
| Atlas purification | Yes | Yes | No | No | Yes | No |

A recent deconvolution algorithm called UXM (Loyfer et al.[1]), does take advantage of methylation haplotypes. Like CelFiE-sum, UXM averages CpGs within a marker region, but unlike CelFie-sum, it does this aggregation within each read individually. Whereas CelFiE has *n* methylation states for *n* cell types, UXM only has two (Table 1, “Num methylation states”=2). One of the two states consists of reads with <25% methylation (the “U” state) and the other consists of reads with >75% methylation (the “M” state). Any read with intermediate methylation is denoted as “X”. The percentage of reads in the U state is calculated for each marker region as U/(X+U+M), or simply the percentage of reads in the unmethylated state. These values are the input for a Non-Negative Least Squares regression to estimate cell type proportions.

Earlier work has used an Expectation-Maximization framework similar to CelFiE to perform methylation haplotype inference [22–25], but only to deconvolute two cell types and not the full *n* cell types which we aim to solve here. We thus sought to adapt the probabilistic features of these earlier methylation haplotype approaches to the problem of *n* cell types.Our first approach to this problem was to conform to the basic CelFiE *n* methylation state model, but use the joint probability of all CpGs within an individual read rather than assigning each CpG site an individual probability (supplementary methods). In order to convey this new aspect, we appended the acronym ISH (Integrated Single-molecule Haplotype) to get *CelFiE-ISH*. We implemented an alternative version of CelFiE-ISH that jointly re-estimates the reference atlas along with the input samples (“ReAtlas” mode), similar to the default algorithm of CelFiE.

Our second methylation haplotype approach adapted the previous 2-cell type methods like MethylPurify[24] and CancerLocator[25] to account for *n* cell types. Similar to those methods, our approach uses two models for each marker region, a low methylation state (or *epistate*) and a high methylation epistate. This is motivated by the finding that epigenetic organization consists of predominantly methylated, predominantly unmethylated, and bimodal regions, with a stochastic noise component [26]. Furthermore, it was recently demonstrated in primary human cell types that cell-type specific methylation manifests primarily as the unmethylated state, with the methylated state being the default background [1]. But unlike UXM, these two states are learned from the data, using methylation haplotypes present across all cell types in the reference atlas. We call this approach Epistate. While both CelFiE-ISH and Epistate use similar Expectation-Maximization models, the differences are substantial. The Epistate method works well only if a low methylation epistate occurs primarily in one target cell type, and if the high methylation epistate is relatively consistent across all other non-target cell types. The *n* state model of CelFiE-ISH does not require this consistency among non-target cell types, but has more free parameters to learn and could thus be more susceptible to overfitting. Like CelFiE-ISH but unlike some earlier implementations of methylation haplotype inference [24], the underlying model of Epistate does not force uniform methylation level across the marker region, but allows discontinuous methylation patterns (one or more highly-methylated CpGs can occur within the low methylation epistate, and vice versa, see Methods for details).

### n-state simulations

We tested the deconvolution of each of the algorithms on simulated data. First, we replicated the simulation model used in the CelFiE paper of Caggiano et al. [21] (Figure 1A). The true methylation proportion of each CpG within each marker region was independently drawn from a uniform distribution for each cell type. For *n* cell types, this would result in *n* different methylation states, each being a Bernoulli process with position-dependent success probabilities. We set the reference atlas depth to 100, approximating the atlas from Loyfer et al. [1]. To examine the effects of read length, we sought to vary the number of CpGs per read from 1 to 10, while keeping the total number of methylation cells (number of reads times number of CpGs) constant. For simplicity, we set the region size to match the read length, and reduced the number of regions with increasing region size to maintain a constant number of CpG sites.

**Figure 1:**
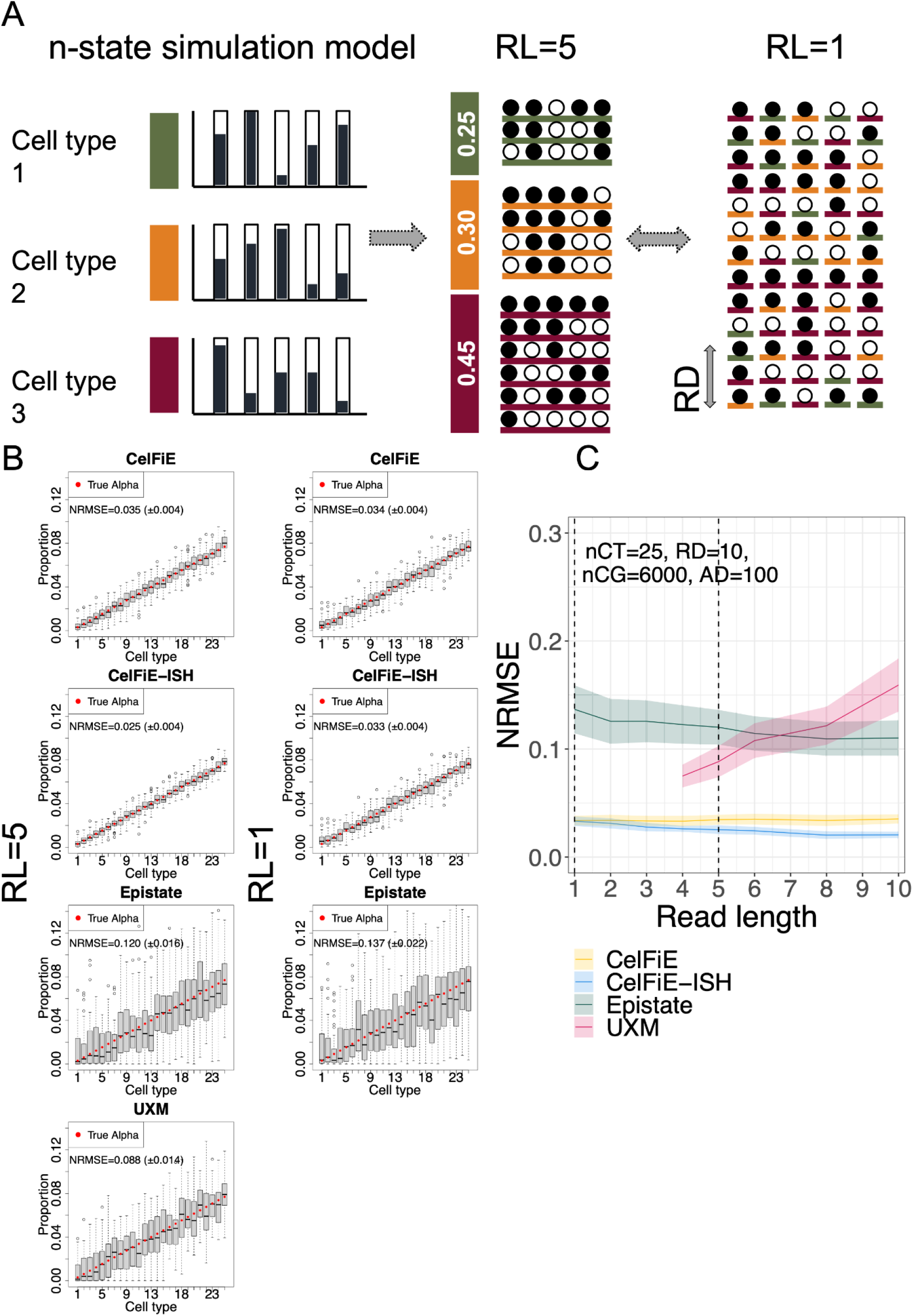
*n*-state simulations. A) schematic drawing of the generative model. The true methylation proportion for each CpG in each cell type is independently drawn from a uniform distribution from [0,1]. Reads are generated from these proportions according to their cell type of origin.Read Length (RL) determines the number of CpGs on each read, and Read Depth (RD) was held constant at 10. B) Estimated proportion of each cell type (gray box) vs. true proportion (red circle) for each model, with read length of 5 (left column) and 1 (right column). Each plot is based on 50 replicate simulations, and shows the Normalized Error Normalized error (Normalized Root Mean Square Error or NMRSE) and NRMSE IQR across replicates. C) NRMSE of each model is shown across a range of read lengths. Shaded areas show standard deviation across 50 replicates. Vertical dotted lines show the condition from panel (B).

We simulated 25 cell types with linearly increasing true proportions, and ran this on each model with varying read lengths (Figure 1B). CelFiE had no difference between read length 1 and read length 5, since it does not use within read methylation haplotypes. This also explains why CelFiE-ISH was nearly identical to CelFiE at rl=1. Summing reduced the variance and limited deconvolution (Figure S1). However, when read length is increased to 5, the improved accuracy of CelFiE-ISH is apparent. These results are summarized in Figure 1C by showing Normalized Root Mean Square Error (NRMSE) for a range of read lengths from 1 to 10. CelFiE and CelFiE-ISH have identical error at rl=1, but CelFiE-ISH decreases in error substantially until rl=4.At the maximum read length of 10, CelFiE had a mean NRMSE of 0.035 (±0.004), while CelFiE-ISH had a mean NRMSE of 0.021 (±0.003), a 40% decrease in error. ReAtlas had no benefit over CelFiE-ISH, although this is likely because the reference atlas did not contain any cross contamination between cell types,which is what ReAtlas is aimed at correcting (Figure S1). It is worth noting that while the overall accuracy is high, some of the methods based on the underlying CelFiE n-state model tend to overestimate the rarest cell types which have proportions close to 0 (Figure 1B). We believe this represents probabilistic uncertainty that does not allow these models to give estimates very close to 0. We will return to this issue in the *in silico* mixture analyses below.

In the 25 cell type simulation, Epistate and UXM performed poorly (Figure 1B-C). Both of these models assume two methylation states, an assumption which is violated in this simulation where each cell type is defined by a unique methylation pattern. We only investigated UXM with read lengths greater than 4, because the UXM algorithm itself removes reads with less than 4 CpGs. Because UXM requires all CpGs in a single fragment to have a consistent methylation state (an assumption violated in this simulation), it performs worse as the read lengths increase. The uniformity of CpGs within a read is not an assumption of Epistate, so while it performs poorly, it does not suffer from poorer performance with increasing read length.

### Two-state simulations

Local consistency of adjacent CpGs in the human genome is a biologically-encoded phenomenon which has been long recognized [27]. For our second simulation condition, we thus simulated reads with local coordination of adjacent CpGs. In this model, we only had two methylation states for each marker region -Θ_*high*_ and Θ_*low*_. Θ_*high*_ is defined as the state with higher mean methylation. Since UXM requires the low state to have average methylation below 0.25, we kept the Θ_*low*_ state at a constant value of 0.1, while varying -Θ_*high*_ between 0.2 and 1.0 (Figure 2A).Each CpG within each read is drawn as a Bernoulli variable with a probability equal to the Θ_*low*_ (for reads derived from the target cell type), and -Θ_*high*_ (for reads drawn from all non-target cell types). The total number of CpGs (nCG=6000) and the number (nCT=25) and proportion of cell types are the same as in the n-state simulations, but we used a greater read depth of 30 reads per cell type (RD=30) since distinguishing two epistates with local coordination is a more challenging problem.

**Figure 2:**
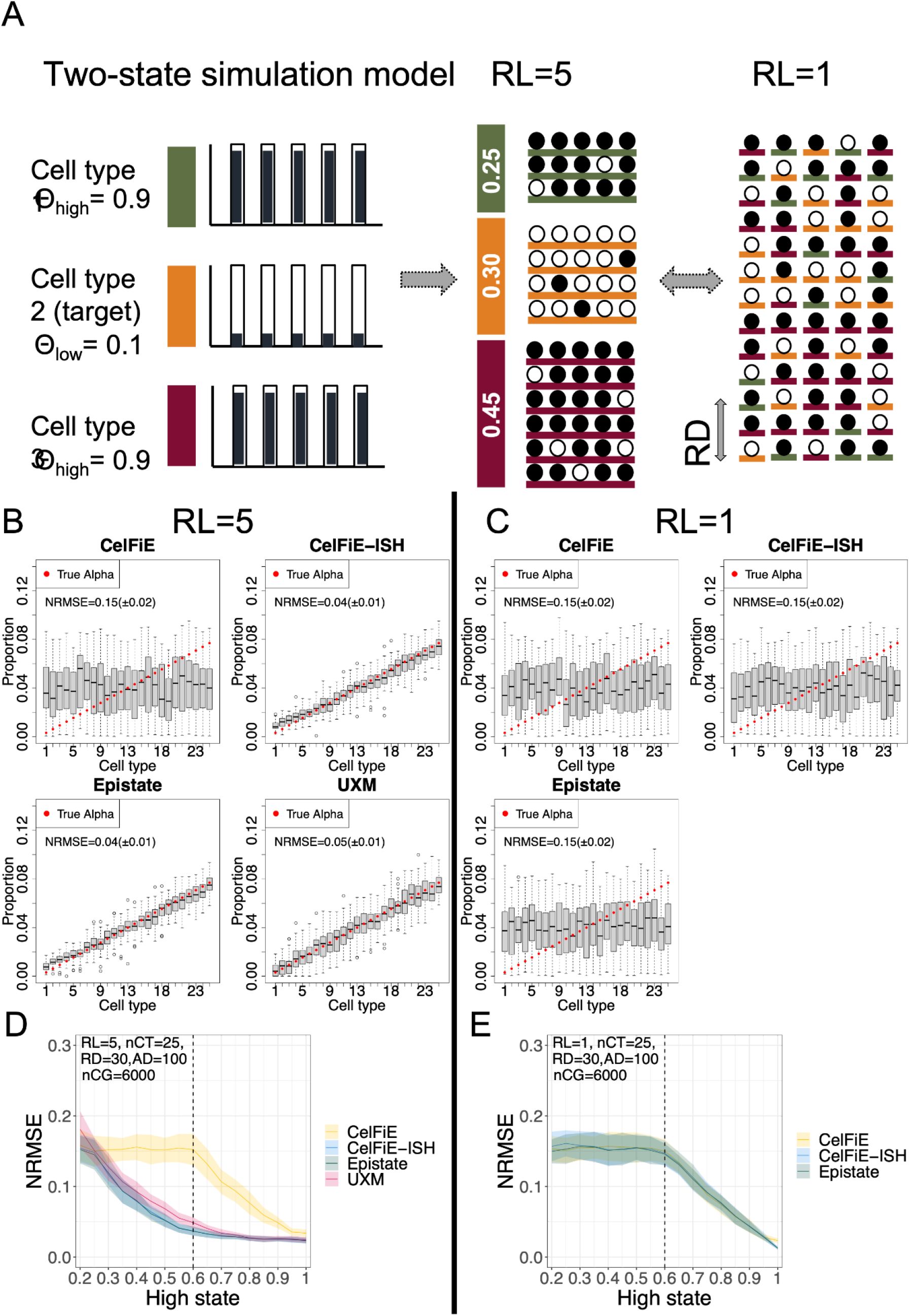
two-state simulations. A) schematic drawing of the generative model. The true methylation proportion for each CpG and cell type is drawn from one of two distributions: low (methylation probability Θ_*low*_ =0.1) and high (methylation probability Θ_*high*_ =0.9). Reads are generated from these proportions according to their cell type of origin.Read Length (RL) determines the number of CpGs on each read, and Read Depth (RD) was held constant at 30. B-C) Simulations using Θ_*low*_ =0.1 and -Θ_*high*_ =0.6, showing the estimated proportion of each cell type (gray box) vs. true proportion (red circle) for each model, using a read length of either RL=5 (panel B) or RL=1 (panel C). Each plot is based on 50 replicates simulations, and shows the Normalized Error Normalized error (Normalized Root Mean Square Error or NMRSE) and NRMSE IQR across replicates. (D-E) NRMSE as a result of varying the -Θ_*high*_ state from 0.2 to 1.0 (with Θ_*low*_ held constant at 0.1. (D) using read length RL=5, and (E) using read length RL=1. Shaded area shows standard deviation across 50 replicates. In (D), a dotted vertical line shows the condition from panel B above, whereas in (E), the dotted vertical line shows the condition for panel C above.

The results of the 25 cell type two-state simulations are shown as Figures 2B-E.With a read length of 1 (RL=1), all models perform similarly - high error rate until the Θ_*high*_ reaches 0.6, and then a linearly decreasing error (Figures 2C,E). With a read length of 5 (RL=5), the basic CelFiE models have exactly the same error profile, which is expected since they can not take advantage of methylation haplotypes (Figures 2B,D). But all of the methylation haplotype aware models (CelFiE-ISH, Epistate, and UXM) have much lower error, with improvement over the basic CelFiE model starting at Θ_*high*_ of 0.3 (which corresponds to only a 0.2 methylation difference between Θ_*low*_ and Θ _*high*_. There is little difference between the haplotype aware models in this simulation.It is worth noting that the two-state simulation model is identical to the Epistate generative model, and complies with all the assumptions of UXM, so it is not unexpected that all models perform well under this simulation. Sum-CelFiE and ReAtlas showed similar patterns to CelFiE and CelFiE-ISH respectively (Figure S2).

### High complexity in silico mixtures

The simulation results demonstrate that the underlying structure of the data has a profound impact on deconvolution. We next tested the models on *in silico* mixtures of real WGBS data, with their true underlying structure. For this we used the deep WGBS data from FACS-sorted cells derived from primary human tissues, which was recently published [1]. Importantly 31 of the 39 cell types profiled had samples from at least 3 individual donors (after removing a small number of samples derived from in vitro culturing, which is known to affect DNA methylation levels. See Methods and Supplemental Table S1). This allowed us to use this resource as both the reference training set as well as the test set using the following hold-out strategy: for each *in silico* mixture trial, we constructed the reference matrix using 2 individual donors, and created the mixture test set from the remaining 1 or more donors. This strategy allowed us to evaluate the models based on individual donors that had never been used in the training data, ensuring that we will not overfit biological aspects of the training dataset.

Marker selection is an important aspect of reference-based deconvolution. We compared several sets of markers for distinguishing the 31 cell types from the Loyfer atlas. First, we called markers using the method provided in the CelFiE package, called Tissue Informative Markers (TIMs) [21]. In the TIM method, CpG sites are selected based on the distance between the percent methylation of a target cell type and the median percent methylation of the CpG across all cell types. Then, a block of +-250bp from the informative CpG is used. We used the default suggested number of TIMs per cell type, 100. As these regions may overlap both within a cell type and between cell types, we merged down the list to non-overlapping genomic intervals. Next, we used the cell-type specific regions identified by Loyfer et al.[1], which included both the top 25 cell type specific markers (the “U25” set), and the top 250 markers (the “U250” set)[1]. The method used to select these markers used blocks of 5 or more CpGs, according to the difference between average methylation of the target cell type against all other cell types.

For our first set of *in silico* mixtures (Figure 3, Figure S3), we used a high complexity mixture, containing all 31 cell types, which were randomly assigned linearly increasing proportions (Figure 3A).Visual inspection shows that at 30x coverage, the TIMs perform relatively poorly compared to the U25 and especially the U250 set (Figure 3A). Since the TIMs markers contain substantially more CpGs than the U25 set (23,913 vs. 7,275, Supplemental Table 1), we attribute this to a superior selection strategy in the Loyfer et al approach, at least for this specific reference dataset. An analysis of error rates across a range of genome coverages (Figure 3B) verifies that error is lowest in the U250 marker set across all samples. In order to ensure that specific cell types were not driving these results, we repeated all benchmarking on two additional random shufflings of cell type proportions (Figures S4-S5).

**Figure 3:**
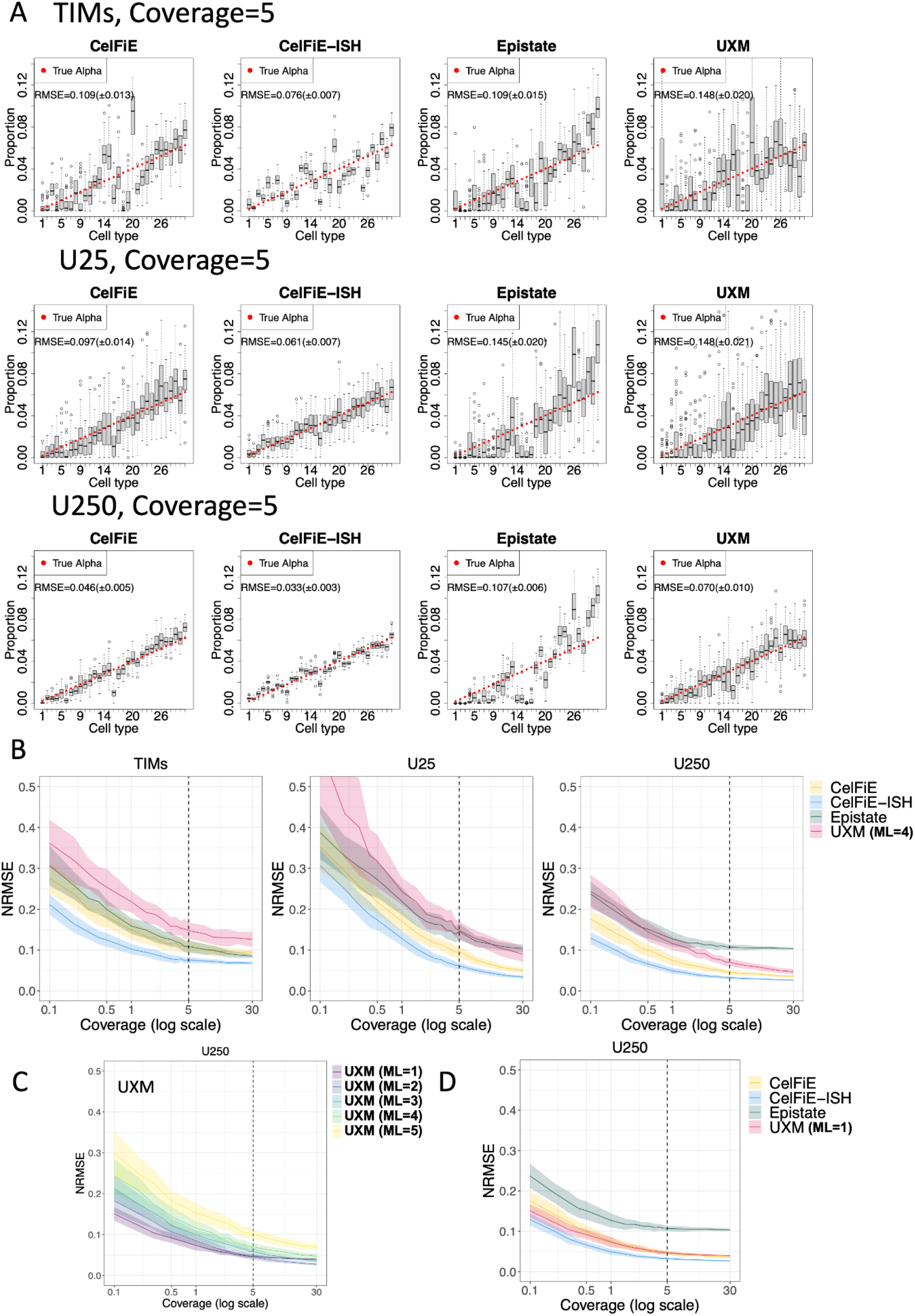
High complexity *in silico* mixtures. 31 cell types from the Loyfer et al. WGBS methylation atlas were mixed at set proportions to yield a total average read depth (coverage) of 5. (A) Deconvolution results using three different sets of input markers, Tissue Informative Marker (TIM) method from Caggiano et al. (top row), Loyfer et al. 25-top markers per cell type U25 (middle row), and Loyfer et al. 250 top markers U250 (bottom row). For each of the four deconvolution models, estimated proportions are shown for each of the 31 cell types as a boxplot of 50 replicates, and true mixture proportions are shown as a red circle. The cell type ordering is listed in the Methods. (B) Mixtures were performed for read depths (coverages) from 0.1x to 30x, showing error (NRMSE) for each model and each marker set. Dotted vertical lines indicate the condition from panel A (coverage=5). Shaded areas show standard deviation across 50 replicates.C) UXM error (NRMSE) for the U250 marker set, at different minimal length (ML) settings. Dotted vertical lines indicate the condition from panel A (coverage=5). Shaded areas show standard deviation across 50 replicates.D) Error (NRMSE) for the U250 marker set, with UXM at a minimal length of 1 CpG. Dotted vertical lines indicate the condition from panel A (coverage=5). Shaded areas show standard deviation across 50 replicates.

Focusing on the U250 set which performed best under all models (Figure 3B, right), CelFiE and CelFiE-ISH had the best performance overall. While performance nearly equalized for CelFiE, CelFiE-ISH, and UXM at 30x coverage, they had substantial differences with lower coverage levels. At 5x, the NRMSE values were 0.046, 0.033, and 0.070, for CelFie, CelFie-ISH, and UXM, respectively. This is a 28% improvement in performance of CelFiE-ISH over CelFiE, and a 53% improvement over UXM.

In the analysis above, we used the default settings for UXM, which include a setting to filter out reads from deconvolution with less than min_length CpGs. Since this is a critical parameter which could remove a substantial portion of the data, and since it was not explored systematically in the original publication [1], we investigated a range of relevant min_length values in our mixture data. Performance increased with decreasing values from 5 to 1, with ml=1 having the best performance, especially at low depths (Figure 3C). When compared to other models UXM with ML=1 had similar performance to CelFiE but still underperformed compared to CelFiE-ISH (Figure 3D). At the 5x level, CelFiE-ISH only had a 30% improvement over UXM ml=1 (NRMSE=0.047), compared to the 53% improvement over UXM default setting, demonstrating the importance of run settings for deconvolution tools. We also looked at alternative versions of CelFiE (Reatlas and sum), but they had very little impact on performance, and in fact ReAtlas performed slightly worse using TIMs (Fig S3). Unlike all other models, Epistate had a systematic bias that overestimated abundant cell types, and this bias was not improved with increasing genome coverage to 30x.

### Circulating DNA - in silico mixtures

To provide a second biologically-inspired mixture scenario and determined the limits of sensitivity, we modeled a cell type background similar to that of cell-free DNA typically circulating in the bloodstream, including lymphocytes (B cells, T cells, NK), myelocytes (monocytes, granulocytes) and hepatocytes. To maximize the available data, we mixed all test sample reads for these 5 cell types (total 100x genome coverage), rather than removing reads to adjust for their relative representation in blood. We then added a “spike-in” of cardiomyocytes reads at varying proportions, and performed full 31 cell type deconvolution using the U250 marker exactly as described above.

Using a total genome coverage of 5x, we compared 50 replicate cardiomyocyte spike-in mixtures to 50 replicates of a null mixture that contained the same blood cell types but no cardiomyocyte reads, and repeated this analysis for three different spike-in proportions: 0.03%, 0.10%, and 0.30% (Figure 4A).Statistically, CelFiE-ISH was better able to distinguish true spike-in vs. null mixtures across the range of spike-in levels, and was the only model to obtain statistical significance down to 0.03%. As with the high complexity mixtures, the CelFiE model variants (Sum and ReAtlas) had little impact on model accuracy (Figure S6). Varying the min_length parameter for UXM produced similar results for ml=2,3,4 and markedly worse performance for ml=1 (Figure S7). The Epistate model performed poorly even at the 0.3% spike-in, as it tends to overestimate abundant cell types and underestimate rare cell types, as discussed above.

**Figure 4:**
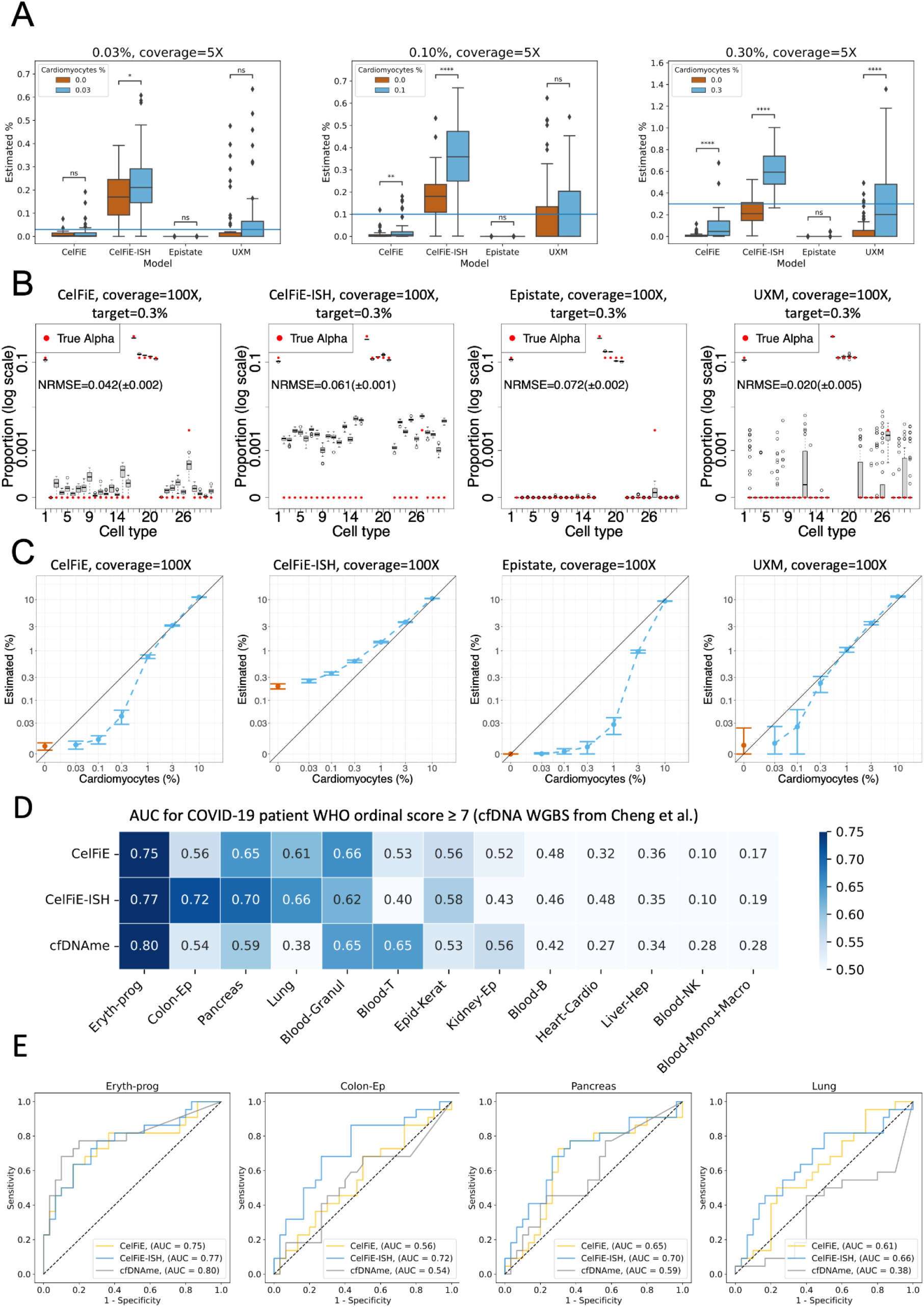
Immune cell spike-in mixtures and circulating DNA in COVID-19 patient plasma. A) Cardiomyocyte samples were added at varying proportions to a background mixture consisting of five cell types commonly found in circulating cell-free DNA: B cells, T cells, NK cells, granulocytes, monocytes, and hepatocytes. The relative immune cell type proportions were held constant while cardiomyocte reads were spiked in at the following percentages: 0.03% (left), 0.1% (middle) and 0.3% (right), using a total genome coverage of 5X. For each model, 31-cell type deconvolution was performed to determine the target cardiomyocyte fraction in 50 replicates of the true mixtures (blue bars), vs. 50 replicates of a null mixture containing no cardiomyocyte reads (red bars). B) Detailed cell type fractions are shown for deconvolutions on a mixture with 100x genome coverage and 0.3% cardiomyocyte spike in. In each case, estimated proportions for each of the 31 cell types are shown as a boxplot of 50 replicates, and true mixture proportions are shown as a red circle. To display the wide range of true proportions, the Y axis is in log scale.The cell type ordering is listed in the Methods. C) 100x genome coverage mixtures were performed with a range of cardiomyocyte spike in levels from 0-10% (blue points), compared to null mixtures with no cardiomyocyte reads (red points). Cardiomyocyte estimates across 50 replicates are shown as standard deviation bars. Both axes are in log-scale. (D) Prediction of COVID-19 severity from WGBS sequencing of 52 patient plasma samples from [28]. Deconvolution from the original paper (cfDNAme) was compared to CelFiE and CelFiE-ISH, after harmonizing the cell types as described in Methods. ROC analysis was performed using the proportions for each cell type separately to predict WHO ordinal code >= 7 (patients requiring mechanical ventilation in the ICU). Cell types are ranked by the maximum AUC across the 3 models. Individual ROC curves for top 4 cell types are shown in (E). A one-tailed t-test was used in (A) to compare the real mixture against the null mixture: * < 0.05, **<0.01, ***<0.001, ****< 0.00001

While CelFiE-ISH was able to best discern true spike in from a null mixture, the estimated cell fraction for the null mixture was significantly elevated above 0 (Figure 4A). To investigate whether this was due to low coverage, we repeated these analyses at the full maximum genomic coverage of our dataset, 100X (Figure 4B-C). CelFiE-ISH values were clearly still elevated, whereas all other models were close to 0 for null mixtures but tended to underestimate proportions for true spike ins at low frequency (Figure 4C). Overall, UXM tended to have the least biased estimates, whereas CelFiE-ISH was the most accurate in distinguishing the rare cell type from the null mixture.

### Circulating DNA - WGBS data from COVID-19 patient plasma

Recently, Cheng et al. [28] reported shallow WGBS sequencing of 52 plasma samples from 28 patients admitted to a McGill hospital for COVID-19 treatment. Of these, 30 were taken from patients in the non-ICU setting (WHO ordinal COVID-19 scores 4-6), and 22 were taken from patients on mechanical ventilation in the ICU (ordinal scores 7-9). Using a methylation-based deconvolution tool, cfDNAme, and a cell type atlas they constructed from publicly available WGBS data, Cheng et al. found that the two strongest predictors of orindal scores >=7 were the total concentration of cfDNA and the concentration of erythroblast-derived cfDNA. Because many cell types had increased cfDNA concentration in the severe patients, we were interested if the relative *proportion* of erythroblast DNA was also associated with severity. Indeed, when we downloaded their original deconvolution, we found that the erythroblast was a much better predictor than any other cell type (Figure 4D and Supplementary Figure S8A).

We performed deconvolution on these samples using CelFiE and CelFiE-ISH (Supplementary Figures S8B-C), and harmonized the cell types from the two atlases to compare AUCs in each shared cell type (Figure 4D-E). Reassuringly, erythrocyte progenitor was the strongest predictor across all 3 methods, and the methods were in generally good agreement for other cell types. However, the next three highest AUC scores (0.72, 0.70, and 0.66) all came from CelFiE-ISH. Colon epithelia and Pancreas were predictive to some extent (AUC>0.5) in all models, whereas lung was only predictive in CelFiE and CelFiE-ISH. In all these cases, CelFiE-ISH yielded a higher AUC than CelFiE, confirming the greater sensitivity observed in the *in silico* mixture experiments above. Lung is particularly interesting, given that COVID is a respiratory disease. While the original study found that the concentration of lung DNA was statistically higher in this COVID cohort than in 4 healthy controls, they did not establish that the proportion of lung DNA was higher, nor did they find any association between lung DNA and severity. While both CelFiE and CelFiE-ISH were able to make this association, the sensitivity of CelFiE-ISH was able to provide a stronger link (AUC 0.66 vs. AUC 0.61).

### Differential marker contributions to deconvolution accuracy

In our *in silico* mixture results, we saw that marker selection (TIM vs. U25 vs. U250) had a major impact on performance of all models. The marker selection approaches used above are relatively ad hoc and none are specifically designed to take advantage of methylation haplotype methods such as CelFiE-ISH and UXM, which should perform better in markers with higher CpG density. We investigated differences between CelFiE and CelfFiE-ISH, which use very similar models but differ in their overall performance and the key aspect of using fragment haplotype information. We adapted our *in silico* mixture methodology to look at the relative contribution and performance of individual marker regions within the U250 marker set. To reduce complexity, we also limited this analysis to a well-characterized set of 6 cell types with pancreas origin. To make the interpretation simpler, we focused on a reduced complexity mixture containing only the 6 cell types derived from pancreatic tissue. This is an ideal set of cell types, because it includes both very distant types (endocrine vs. epithelial vs. endothelial), as well as several highly related cell types (alpha vs. beta. vs. delta).

To facilitate this analysis, we used a simplified version of the Expectation-Maximization model to quantify the contribution of each marker independently. We start with the full set of reads derived from a holdout donor/sample of the target cell type (for instance the top group in Figure 5A is colored yellow and is a sample of pure pancreatic Delta cell reads). We then run a simplified version of the EM deconvolution, to estimate the proportion of each of the 6 cell types for this pure population of reads. The standard EM algorithm starts with a random prior for cell type proportions and adjusts these until convergence. In this simplified version no prior is employed, and therefore only a single E-step and a single M-step are performed. A “perfect” marker would give a proportion of 100% to the target cell type and 0% to all other cell types. We performed this on each of the U250 markers from each of the six pancreatic cell types, and sorted markers by how close to perfect they were (Figure 5A). Results for the TIM regions and U25 marker sets are shown in Figure S9 and S10 respectively.

**Figure 5:**
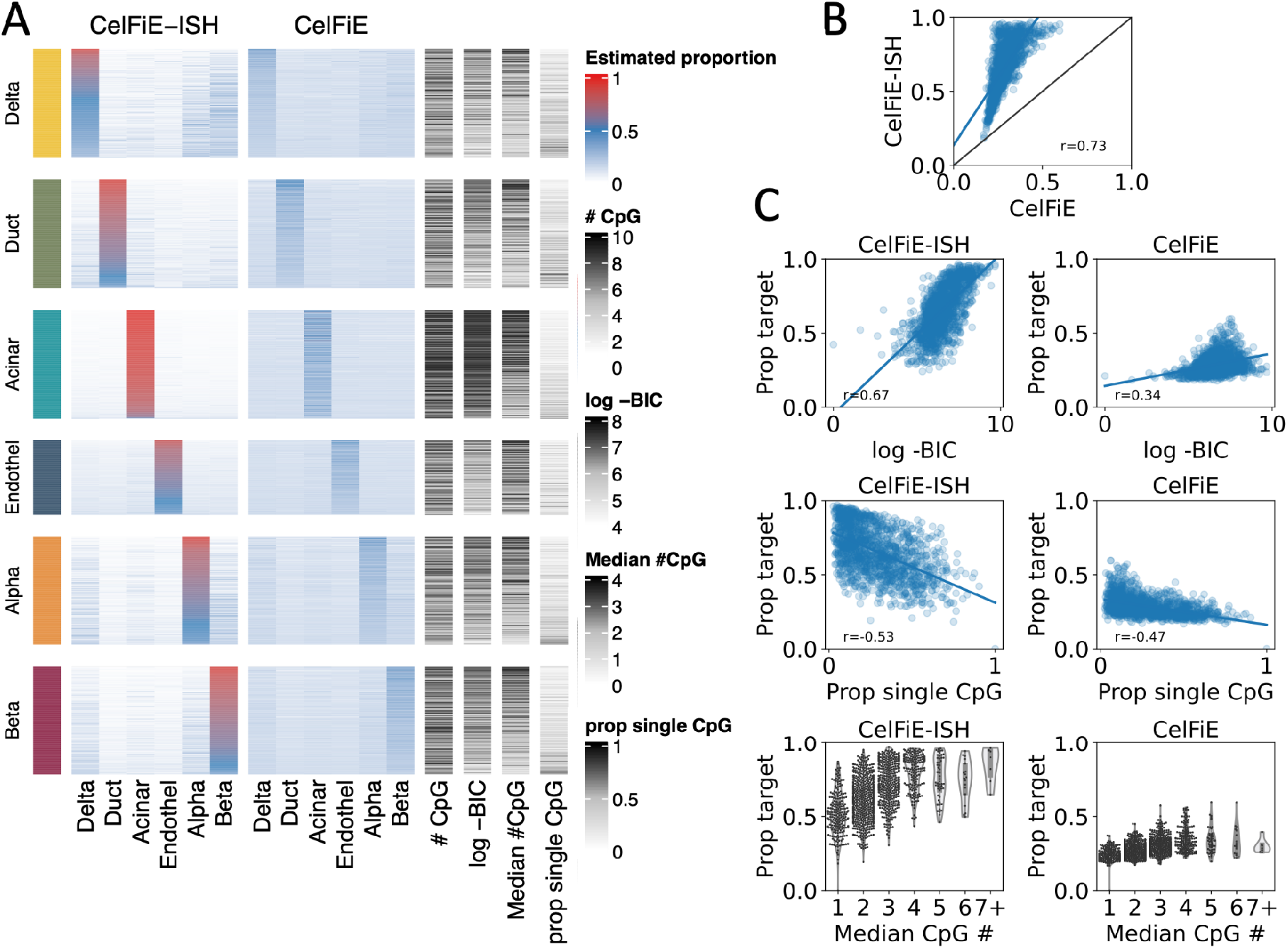
Differential marker contributions to deconvolution accuracy. A) Performance of individual U250 markers in identifying the cell-type specificity between 6 pancreatic cell types. Each group on the horizontal axis shows the accuracy of identifying reads from a specific cell type (i.e. the “yellow” section is performance only on reads derived from the pancreatic Delta cell holdout sample). Within each group all U250 markers are displayed as rows, and the percentage of reads identified as each of the 6 cell types in a 6-cell type deconvolution is shown as a heatmap. Rows are ordered by accuracy, i.e. markers that assign the highest fraction of reads to the target cell type are at the top. Additional features of each marker are shown at the right: # CpG: number of CpG sites in the region, log -BIC: log of the Bayesian information criterion, Median # CpG: the median number of CpGs per read, prop single CpG: proportion of reads with a single CpG out of all reads overlapping the region. B) Correlation of the percentage of reads assigned to the target cell type (“Prop target”) between CelFIE and CelFiE-ISH. C) Correlation of the percentage of reads assigned to the target cell type (“Prop target”) between each model and each individual marker features (left column for CelFIE-ISH, right for CelFiE).

Overall, CelFiE-ISH performed better than CelFiE, as indicated by the higher values along the diagonal of the heatmaps in Figure 5A. High values off the diagonal indicated confusion patterns among the cell types, and these appear to be similar between the two models, for instance significant confusion Delta, Alpha, and Beta cells. If we make a scatter plot of the on target percentages (the values in the diagonal), we can see that CelFiE and CelFiE-ISH are well correlated, but CelfiE-ISH is universally more accurate (Figure 5B).

We next sought to determine which marker features predict high performance in this analysis. We characterized several features that we would expect to influence methylation haplotypes and thus favor the haplotype-based CelFiE-ISH model - these features included the proportion of reads with only a single CpG (“Prop single CpG”), and median number of CpGs per read (“Median CpG #”), and a measurement of the bimodal-ness of reads derived from the EM model (Bayesian Information Criterion or “BIC”, see Methods). As expected, all of these features were more strongly correlated with CelFiE-ISH than with CelFiE (Figure 5C). The feature directly related to bimodal-ness of read haplotypes (BIC) was much more strongly correlated in CelFiE-ISH vs. CelFiE (p=0.67 vs. p=0.34), whereas those related to CpG density were much more similar in the two models. The correlation of these features in CelFiE suggests that high CpG density is a strong predictor of marker performance overall, not just in haplotype models, and should thus be considered important for in all future marker selection efforts for whole-genome DNA methylation-based deconvolution.

## Discussion

Methylation arrays and methylation sequencing are both commonly used for profiling of clinical patient samples. Methylation arrays provide information at a single base level, while methylation sequencing can provide within-read methylation patterns or *methylation haplotypes* [22]. A single read can only have one cell type of origin, and a recent whole-genome atlas of purified cell types [1,29] allows for general purpose multi cell-type deconvolution. Deconvolution methods developed for methylation array data [7,19] do not take methylation haplotype information into account. A new read-aware method, “UXM”, was published along with the methylation atlas [1,29]. UXM relies on percent methylation within each read, and thus forgoes base-level information. Here, we explore two new read-aware algorithms, that utilize both the position across the genome and within-read information. One of these (CelFiE-ISH) is an extension of the model in CelFiE [21], which is the leading non-read aware algorithm. We did not evaluate other methods that were based on array [7], or not general with respect to cell-types [8,20,24,30] or whole-genome markers [11].

First, we generated purely artificial WGBS data and varied the number of CpG sites on each molecule. In one simulation model where each cell type had a different methylation state (n-state model), the number of CpGs sites per molecular only had a modest effect the difference between the read-aware approach (CelFiE-ISH) and the non read-aware version (CelFiE). In a second model where there are only two primary methylation states (2-state model) and one state was unique to a single cell type, more CpGs per read made a dramatic improvement on the read-aware CelFiE-ISH.

For more realistic simulation experiments, we mixed reads from individual human cell types from the human WGBS methylation atlas [1], carefully creating hold-out test datasets that contained no samples from the same individual donors used in the training dataset. Here, CelFiE-ISH had a significant advantage over CelFiE, as well as UXM, but only about 30% improvement, not nearly as strong as seen in the 2-state simulation model. While this improvement may appear modest, the advantage was particularly apparent for rare cell types - unlike CelFiE or UXM, CelFiE-ISH was able to detect a cell type present in just 0.03% of reads out of a total of 5x genomic sequencing coverage. In a completely independent WGBS dataset of COVID-19 patients, this allowed CelFiE-ISH to find stronger associations with rare cell types such as lung epithelium. While CelFiE-ISH performed best at statistically distinguishing rare from non-existent cell types, the *in silico* mixtures revealed an overestimation of both. One possible strategy to mitigate this behavior would be to implement weighting of individual reads [31]. Long reads would be assigned bigger weights and short, ambiguous reads would be down-weighted.

The second algorithm we produced, Epistate, was based on the two-state simulation model. Even in data produced from two-state simulations, it performed no better than CelFiE-ISH. It also suffered from a strong bias for more frequent cell types which limited its performance in more realistic *in silico* mixture experiments. It is possible that our model is overfitting the reference data by trying to estimate cell type proportions in the reference atlas itself - future work should investigate this.

The final read-aware algorithm, UXM, performed somewhat poorly using the default parameters, but when we expanded the search for parameters, performance could be improved by decreasing the minimum number of CpGs per read used (min_length or ML) was from 4 to 1. However, setting ML to 1 had a detrimental effect on the second mixture scenario where the number of reads from the target cell type was small. This shows that the optimal UXM settings might be dependent on read depth and/or target cell type proportions. CelFiE and CelFiE-ISH do not require such settings since they are probabilistic and incorporate the number of CpGs on the read implicitly.

Our study also revealed the strong dependence on the type and number of cell-type specific marker regions used. The difference between markers used by the CelFiE package (TIMs, [21]) and the markers used by UXM (U250) [1] was just as important as the deconvolution algorithm. Both marker selection strategies try to capture regions with low intra-cell-type variance and high inter-cell-type methylation difference, but they use different heuristics and relatively arbitrary cutoffs (the best *n* markers per cell type where *n* is an arbitrary number, etc.) Both the TIMs and the U25/250 regions were selected based on beta values, though the TIM method considers single CpG sites and the Loyfer method considers blocks of CpGs. A recent novel approach to marker selection employs three strategies: one-tissue-vs-the-rest as in the Loyfer and TIM markers, one-group-vs-another-group based on phylogeny, and one-tissue-vs-another-tissue to distinguish highly similar cell types [11].

We attempted to shed light on the selection of marker regions by evaluating performance of individual markers within the U250 set, based on accuracy in our *in silico* mixture train/test datasets. We found that there was a large variability of performance across these markers. Under both models, performance was associated with higher CpG density, a feature which should be considered explicitly when selecting markers for any deconvolution method. A measure of the “bimodal-ness” of the methylation haplotypes across reads (BIC between a two-epistate model and a one-epistate model) was strongly associated with marker performance in the read-aware CelFiE-ISH model but much less so in the CelFiE model, indicating that markers should be tailored to the deconvolution approach being developed. Features specific for methylation haplotypes could be used, such as the average number of CpGs per read. Alternatively, a train/test strategy could be used to evaluate marker regions as we did above, subject to the challenges of designing hold-out test datasets that are truly independent of the training data. Validation using independent datasets is important, especially before using the markers to design fixed capture panels for future studies [1,10]. In the current sequencing landscape, markers may also require tailoring for different sequencing platforms. Sequencing technologies from Oxford Nanopore and Pacific Biosciences sequence DNA methylation natively from tissues with fragments in the 5-20kb range [32], which should translate to higher accuracy for read-aware approaches like CelFiE-ISH if longer markers are selected.

## Methods

Notations are consistent with Caggiano et al. [21]. We are provided with a reference atlas composed of *T* pure cell types indexed by *t*, at *M* CpG sites indexed by *m*. We only look at marker regions,i.e. regions with methylation differences between cell types, assuming each read *c* spans no more than one marker.

The reference atlas consists of one matrix *β*_*t,m*_, with the probability of methylation for cell type *t* at position *m*. We receive a mixture, *X*, composed of *C* reads. We assume each WGBS read is drawn from some cell type *t* at some marker *m*.

*Z*_*t,c*=_1 indicates that is the cell-type of origin for read *c*.

*We assume the mixture is drawn from a multinomial distribution*:

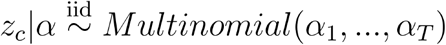

*Where α*_*t*_ *is the probability that a read originates from cell-type t*.

### The CelFiE-ISH algorithm

The mixture retains within-read information, and is represented by one matrix *X*_*c,m*_, with dimensions *C* reads over *M* CpG sites.

This algorithm does not re-estimate the reference atlas. The complete data likelihood is:

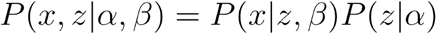

In the E-step, we use the joint probability of an entire read:

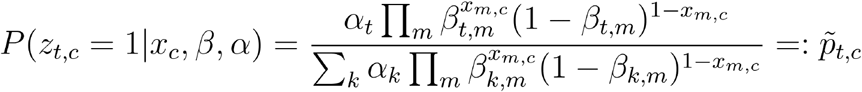

The M-step is equivalent to the first part in the CelFiE algorithm:

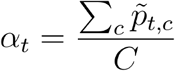

The full derivation of the algorithms is available in the Supplementary Note.

### The CelFiE-ISH ReAtlas algorithm

As in CelFIE, the reference atlas consists of two matrices, *Y*_*t,m* and_ 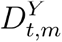, with the number of methylated and total reads for cell type *t* at position respectively. We assume *Y*_*t,m*_ is drawn from a Binomial distribution with *β*_*t,m*_ being the true methylation probability and 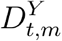 being the number of trials. The complete data likelihood is:

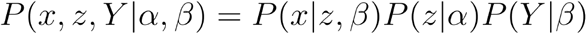

We re-estimate the atlas at each M-step:

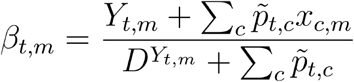

### The Epistate algorithm

At every marker region, reads are drawn from one of two possible states: *θ*_*low*_ and *θ*_*high*_. Each state consists of a set of binomial distributions θ_=_{ *θ*_1,_ *θ*_2,*…*_ *θ*_*m*_}, one per CpG site covered by the marker region. *θ*_*high*_ is arbitrarily defined to be the epistate with higher mean methylation.Cell types differ by the probability of observing each epistate in each region, denoted by λ.

The reference atlas consists of λ, *θ*_*high*_ and *θ*_*low*_. They are estimated separately with their own EM model. *μ*_*high,c*_ = 1 indicates that read belongs to Θ_*high*_. The complete data likelihood is:

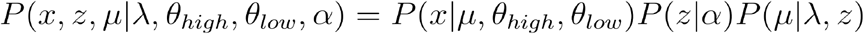

In the E-step, we estimate both *z* and *μ*:

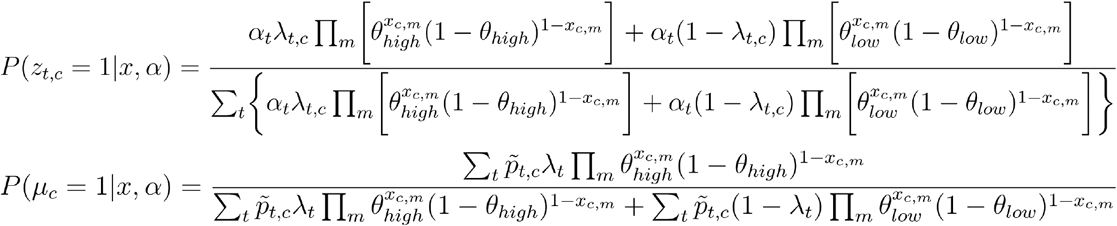

The M-step is the same as CelFiE-ISH.

### Other deconvolution methods

CelFiE was run using code available at https://github.com/christacaggiano/celfie. UXM deconvolution was run using code available at https://github.com/nloyfer/UXM_deconv. We created a custom script to calculate %U reads from epiread files. The U threshold was 25% and M threshold was 75%. Only reads with 4 or more CpGs were considered, for both the reference atlas and the input samples.

### Computational cost

CelFiE scales linearly with the size of input and reference. For a single individual, it is O(T, M). All the read-based models iterate over each read C, making CelFiE-ISH, CelFiE-ISH ReAtlas and Epistate O(C,T,M). This means the read-based models scale linearly with the number of reads, cell types and CpG sites. Generating a reference atlas of methylated calls and total calls for CelFIE, CelFiE-ISH and CelFiE-ISH ReAtlas requires summing over each position in the genome for each reference cell type, which is O(C, T, M). Epistate requires estimating λ, *θ*_*high*_ and *θ*_*low*_ for the atlas. This is done in a read-based approach in O(C,T,M).

### n-state simulations

In this simulation, a cell type was defined as a Bernoulli process. The probability of methylation at each position was drawn from a random uniform distribution between 0 and 1. To simulate a read mixture of depth *RD*, from *T* cell types with proportions, we first calculated the total number of reads in the mixture, taking into account the read length (*RL*):

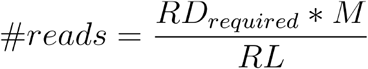

We then sampled *α*_*t*_ to receive the number of reads per cell type. Next, the methylation status of each CpG in each read was determined by binomial sampling of its cell type of origin, at the appropriate position, *β*_*t,m*_. Unlike the simulations in Caggiano et al.[21] and Keukeleire et al.[29], the read depth was a set parameter and not sampled. The total number of individual methylation cells (*RD* * *M*) was held constant regardless of read lengths by adjusting the number of reads. This is essentially equivalent to dividing the CpG sites to blocks/regions of size *RL*.

To transform the atlas to a format compatible with Epistate+, we set *θ*_*low*_ to be *β*_*t,m*_ of the target cell type, and *θ*_*high*_ to be mean *β*_*t,m*_ of all other cell types. For CelFiE, we multiplied *β*_*t,m*_ by the atlas depth to receive methylation and coverage data.

Root mean squared error (RMSE) was normalized to the worst possible RMSE, equivalent to estimating the cell type with the lowest true proportion as the sole contributor:

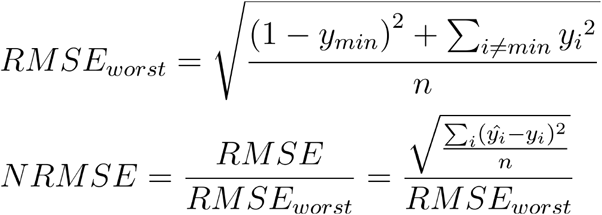

### Two-state simulations

As in the Epistate model, this simulation includes only two methylation states: *θ*_*low*_ and *θ*_*high*_. Therefore, to deconvolute more than two cell types, more than one region is required. At each region, the probability of methylation for the target cell type was set to *θ*_*low*_ and all other cell types were set to *θ*_*high*_. While not all the models require it, in this simulation we kept *θ*_*low*_ and *θ*_*high*_ constant at all CpG sites in the region.. *θ*_*low*_ was set to 0.1, and *θ*_*high*_ varied, but was always >0.1. We kept *λ*_*t*_ binary, so that *θ*_*low*_ was exclusively associated with the target cell type. We explore non-binary *λ*_*t*_ values in Figure S11.

To create a mixture, we used the same process described above: calculated the number of reads for each cell type and sampled the methylation states accordingly. Likewise, the same adjustments for read length were made, and the total number of methylation cells was kept constant.

To generate an atlas compatible with all models except Epistate, we calculated the expected beta value:

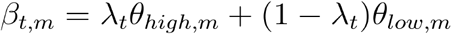

### WGBS data processing

BAM files for the reference atlas were retrieved from Loyfer et al.[1] at European Genome-phenome Archive (EGA) under study accession number: EGAS00001006791 and re-processed with the nf-core/methylseq pipeline [33,34] using the biscuit aligner (https://github.com/ekushele/methylseq/blob/master/main_bams.nf). Briefly, the quality of fastQ files was assessed using FastQC (v0.11.9). TrimGalore (v0.6.4_dev) was used to trim adapters from all samples with the following parameters: ‘clip_r1 = 10 clip_r2 = 15 three_prime_clip_r1 = 10 three_prime_clip_r2 = 10’. We aligned the trimmed samples to the hg19 genome assembly, using the biscuit aligner (v0.3.15.20200318), with ‘-b 1’ for directional library. Duplicates were marked using samblaster (v0.1.24) with ‘--addMateTags’. Then, BAM files were sorted and indexed using Samtools (v1.9). The sorted and indexed BAMs were given as an input to ‘biscuit pileup’, to generate VCF files. Bed files were generated from vcf using ‘biscuit vcf2bed’ with ‘-k 1 -t cg’ arguments. We implemented a custom approach to filtering SNPs and generating Biscuit epiread files which contained a merged version of the two ends of each paired-end read. The code is available at https://github.com/ekushele/methylseq, and details are given in the Supplementary Note. Since native Epiread files contain SNP genotypes, we masked all SNP genotypes and provide these unprotected files at GEO GSE239605.

### Marker regions

To select TIMs, we used the code from Caggiano et al.[21], with a minimal depth of 15, 1 missing value, and 100 regions per cell type. We used blocks of +=250bp around each CpG. For 31 cell types, 2617 TIMs were found. These were merged down to 1755 non-overlapping regions, using an interval union operation for each overlapping set of regions. Regions that were included for more than one cell type were removed, resulting in 1665 regions unique to a single cell type. We used only autosomes for the analysis.

We obtained the Loyfer U25 and U250 marker region lists from Supplemental Tables S4A and S4B from [1], and removed markers associated with the excluded cell types, as well asregions within sex chromosomes. This resulted in 770 total markers for U25 and 7475 total markers for U250. All regions used are available in table S2.

### In silico mixtures - High complexity mixtures

For the *in silico* experiments, we used hold-out samples, and created an atlas from all other samples. For the pan-atlas analysis, we started with the Loyfer 205 sample atlas. We excluded 15 samples that were cultured *in-vitro*, since this can affect methylation levels.

We filtered the remaining samples for groups with at least 3 biological replicates. This left 31/39 groups, and 185/205 samples. For each simulation, we held out one sample (derived from a single donor), and created the atlas from the other 2 or more samples (which had different donors than the hold-out sample). In some cases, multiple cell types were derived from a single donor - in this case, we always used the same donor for the hold-out samples, so that no samples from this donor would ever be included in the reference atlas.Complete details on each sample are provided in the supplementary Table 1.

For the n=31 cell types in the high-complexity mixture, we calculated the cell type proportion vector as 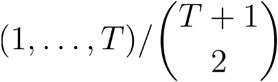 with T=31 [21].

We assigned each *t* a random cell type. In the simulation used in the main figures, this ordering of t=1 to T=31 is as follows: Blood-B, Eryth-prog, Gastric-Ep, Breast-Luminal-Ep, Pancreas-Alpha, Ovary+Endom-Ep, Small-Int-Ep, Lung-Ep-Alveo, Adipocytes,

Breast-Basal-Ep, Prostate-Ep, Oligodend, Pancreas-Duct, Head-Neck-Ep, Endothelium, Neuron, Blood-T, Blood-NK, Blood-Granul, Blood-Mono+Macro, Liver-Hep,

Pancreas-Acinar, Bladder-Ep, Thyroid-Ep, Kidney-Ep, Pancreas-Delta, Heart-Cardio, Colon-Ep, Lung-Ep-Bron, Pancreas-Beta, Fallopian-Ep. Additional shufflings of the cell-type order are shown in figures S4-5.

To sample the mixture to a predefined read depth (RD), we calculated the total number of reads to sample across all marker regions, and then sampled from all reads that overlapped marker regions including partial overlaps, according to the cell type proportions given above. The number of reads for a given cell type was sampled across all marker regions together, so individual marker regions can deviate slightly from the global read depth due to sampling.

The required number of reads is the desired read depth (RD_required) times the total number of CpG sites in the marker regions (M), divided by the average read length RL (CpGs per read). This is equivalent to the required read depth times the total number of reads in the marker regions (C) divided by the read depth in the mixture samples (RD_input).

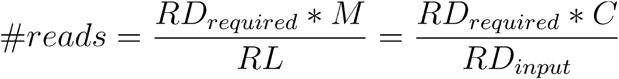

The same random mixture files were used as input for all deconvolution models.

### In silico mixtures - Circulating DNA inspired mixture

The cfDNA-like background was composed of Blood-B, Blood-T, Blood-NK, Blood-Granul, Blood-Mono+Macro and Liver-Hep. As in the high complexity mixtures, we used separate samples and donors for the reference and mixtures. The hold-out samples used to create the mixture included 3 T cell samples and one of each other cell type (supplementary Table 1). To maximize the possible coverage, the background consisted of 37.5% T cells and 12.5% of each remaining cell type.We added a spike-on of cardiomyocytes at varying proportions, reducing the background accordingly. For example, the 3% cardiomyocyte mixture included 36.375% T cells and 12.125% of each remaining cell type, summing to 97% background.

We sampled the mixtures in the same process described above for the high complexity mixtures.While only a few cell types were used to create the mixture (had *α*_*t*_ >0), the full 31 group atlas was used for deconvolution. Cell type ordering is identical to the order in the high complexity mixtures detailed above.

### COVID-19 cfDNA analysis

Publicly available FASTQ files for 52 WGBS plasma samples from Cheng et al. [28] were downloaded from the Short Read Archive (SRA) Project PRJNA687910. The original cfDNAme cell-type deconvolutions and metadata for these samples were downloaded from Supplemental Table 1 of ref [28]. For analysis with CelFiE and CelFiE-ISH, the FASTQ files were processed to methylation Pat files using wgbs_tools (https://github.com/nloyfer/wgbs_tools) as described in [1].

Because this is an independent dataset, we did not use the 31 cell type atlas used above as a holdout training dataset. We constructed a reference atlas containing all 39 cell types described in Loyfer et al. [1], using all 205 samples (table S1), on U250 regions, excluding sex chromosomes, retrieved from Loyfer et al. supplementary table 4B [1]. Beta values for these regions are available at https://github.com/methylgrammarlab/deconvolution_models. We performed deconvolution using the full 39 cell types. Based on these results, we performed ROC analysis in two different ways. First, we used all 39 cell types for the CelFiE and CelFiE-ISH models (shown in Supplementary Figure S8). For direct comparison of the different methods (shown in Figure 4), we harmonized the cell type atlases as follows. We merged more granular cell types in the HUJI atlas into more general ones from [28] by summing cell type proportions, as follows: All pancreatic cell types (Pancreas-Alpha, Pancreas-Beta, Pancreas-Delta, Pancreas-Acinar and Pancreas-Duct) were summed to a composite “Pancreas” cell type and Lung-Ep-Bron and Lung-Ep-Alveo were summed to a “Lung” cell type. Similarly, cfDNAme “monocyte” and “macrophage” were summed.

1-to-1 cell type mappings for other cell types were as follows (first is cfDNAme, second is CelFiE/CelFie-ISH): ‘monocyte’+’macrophage’: ‘Blood-Mono+Macro’, ‘neutrophil’: ‘Blood-Granul’, ‘Tcell’: ‘Blood-T’, ‘Nkcell’: ‘Blood-NK’, ‘Bcell’: ‘Blood-B’, ‘erythroblast’: ‘Eryth-prog’, ‘kidney’: ‘Kidney-Ep’, ‘heart’: ‘Heart-Cardio’, ‘skin’: ‘Epid-Kerat’, ‘lung’: ‘Lung’, ‘liver’: ‘Liver-Hep’, ‘pancreas’: ‘Pancreas’, ‘colon’: ‘Colon-Ep’.

We performed ROC analysis as follows. 30 “Negative” samples were defined using the Cheng et al. metadata fields timepoint_ordinal between 4-6, and 22 “Positive” samples were defined as timepoint_ordinal 7-9. True positive rates, false positive rates, and AUC were calculated using *sklearn*.*metrics*. Some cell types corresponded to AUC scores less than 0.5 (anti-correlated with severity).

### Informative region ranking

For informative region ranking, we used only the pancreatic cell types: Pancreas-Beta, Pancreas-Deta, Pancreas-Duct, Pancreas-Alpha, Pancreas-Acinar and Endothelium derived from pancreas samples. Each of these had at least 3 individual donors.We employed a leave-one-out approach, using all samples except one individual donor for the reference atlas, and the other donor for testing. For this analysis, we generated a downsampled “mixture” that is 100% the target cell type. Then we performed a deconvolution for each marker region individually. We performed this analysis several times, using each individual donor as the holdout sample in one trial. We then averaged the estimated proportions across all individuals (weighted by the number of reads derived from each).

All of the deconvolution models are designed to sum over all marker regions. We adapted them to assign an output to individual marker regions, as follows. We removed the prior from the E-step for each model, i.e. the *α* parameter. For CelFiE and sum-CelFiE that would be:

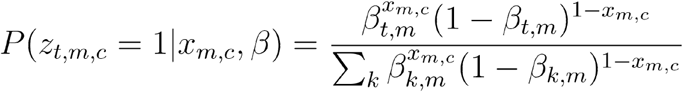

For CelFiE-ISH and CelFiE-ISH ReAtlas:

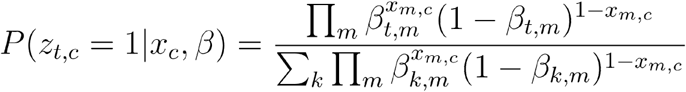

For Epistate:

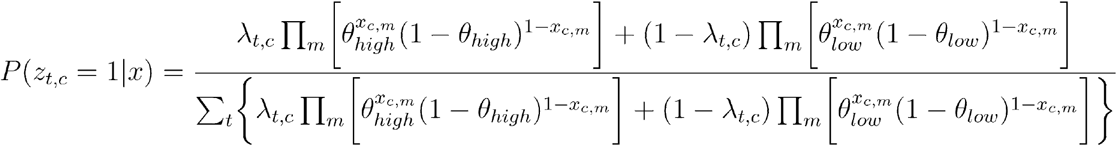

Statistics were calculated across all samples jointly: number of CpG sites in the region, the median number of CpGs per read and proportion of reads with a single CpG out of all reads overlapping the region (including partial overlaps). To assess bimodality, we employed the Bayesian Information Criterion (BIC) to compare the likelihoods of a one-state model and a two-state model. We adapted the two-state model described by Fang et al. [23] to accommodate any mixing ratio (supplementary methods), and compared its likelihood to the one-state model.

## Author contributions

The authors confirm contribution to the paper as follows: study conception and design: B.P.B.,I.U.; analysis and interpretation of results: I.U., E.K, T.J.T, D.A.; draft manuscript preparation:I.U., B.P.B. manuscript editing: I.U, B.G., E.K., B.P.B. All authors reviewed the results and approved the final version of the manuscript.

## Declarations

BPB is a consultant for VolitionRx.

## Funding

BPB received support from the Israel Science Foundation personal grant 2279/22, as well as startup support from Hebrew University, the Kamea B program of the Israel Ministry of Aliyah and Immigrant Integration, and the Beethoven Foundation. IU and EK were funded by a Council for Higher Education grant to the Hebrew University Center for Interdisciplinary Data Science Research (CIDR) to BPB and BG. IU was also supported by the Azrieli foundation.

### Acknowledgements

We thank Tommy Kaplan and Netanel Loyfer for providing important feedback on our manuscript, as well as advising us on use of datasets from the HUJI WGBS atlas (*Nature* 2023, PMID 36599988). We thank Netanel Loyfer for processing the COVID-19 samples to pat files. We also thank Yuval Dor and Amir Eden for helpful discussions and advice. Computation was performed on the Hebrew University Research Computing Services cluster (HURCS), with assistance from Yaron Weitz and Ori Adam.

## Data availability

For WGBS data used to build models and perform in silico mixtures, raw sequence data were taken from the Hebrew University human cell type methylation atlas, available European Genome-phenome Archive (EGA) under study accession number: EGAS00001006791, and processed as described in the Methods. We have deposited processed data files at GEO accession GSE239605, including epiread files with all SNP information masked. For validation WGBS data for COVID-19 patients, publicly available sequence FASTQ files were downloaded from the Short Read Archive (SRA) Project PRJNA687910 and processed data files were obtained from [1] (GEO GSE186458).

## Code availability

CelFiE-ISH and all other deconvolution models are available at https://github.com/methylgrammarlab/deconvolution_models. Code to reproduce simulated datasets and in silico mixed datasets are available from https://github.com/methylgrammarlab/deconvolution_simulation_pipeline and https://github.com/methylgrammarlab/deconvolution_in_silico_pipeline, respectively. All source code is published under the permissive MIT open source license.

## Supplementary Information

**Figure S1:**
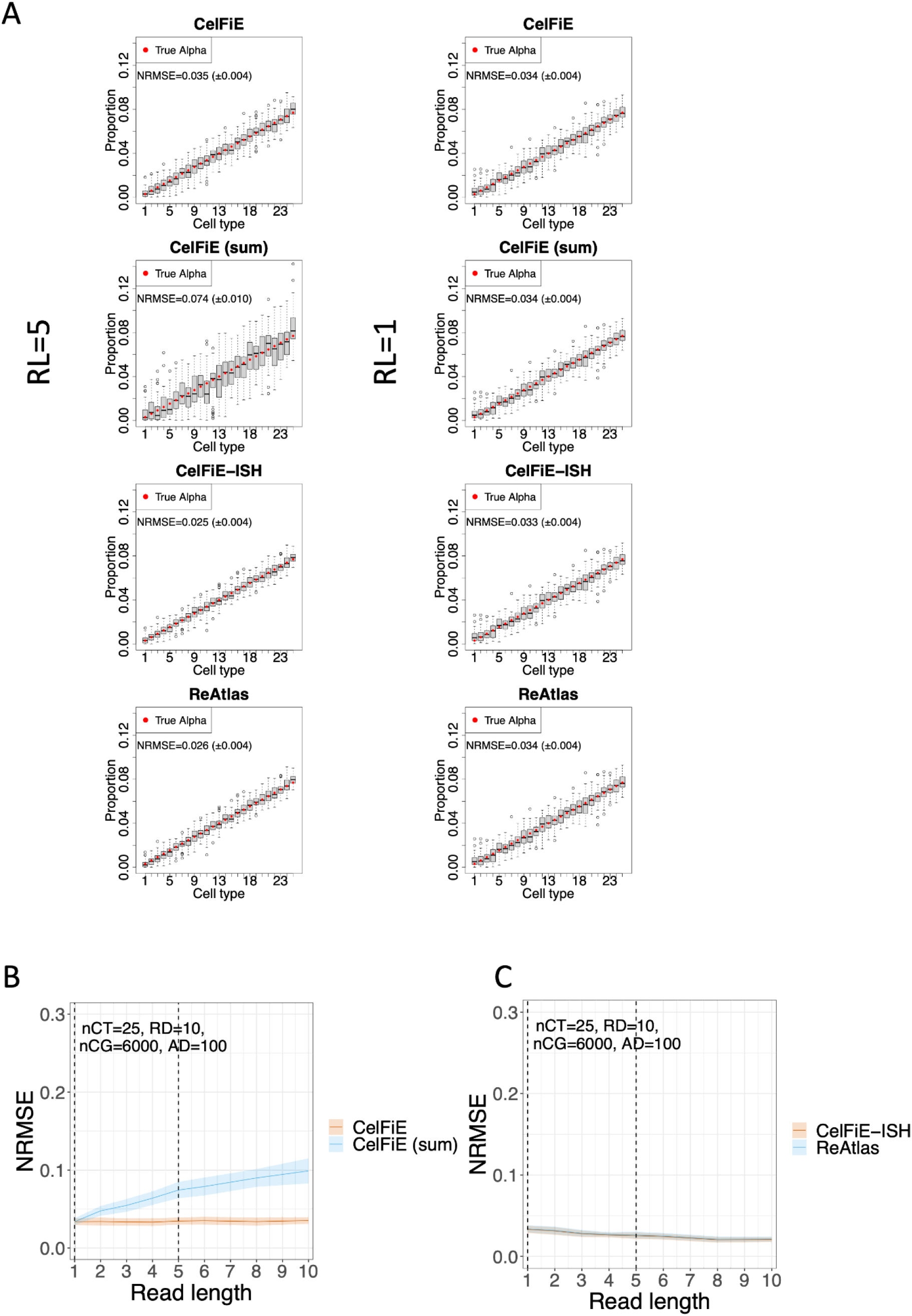
n-state simulations. A) Estimated proportion of each cell type (gray box) vs. true proportion (red circle) for each model, with read length of 5 (left column) and 1 (right column). Each plot is based on 50 replicate simulations, and shows the Normalized Error Normalized error (Normalized Root Mean Square Error or NMRSE) and NRMSE IQR across replicates. B, C) NRMSE of each model is shown across a range of read lengths. Shaded areas show standard deviation across 50 replicates. Vertical dotted lines show the condition from panel (A). CeFiE-Sum compared to CelFiE (B) and ReAtlas is compared to CelFiE-ISH (C). Since we kept the region size the same as the read length, with increasing region size the summed version of CelFiE loses information and performs poorly.

**Figure S2:**
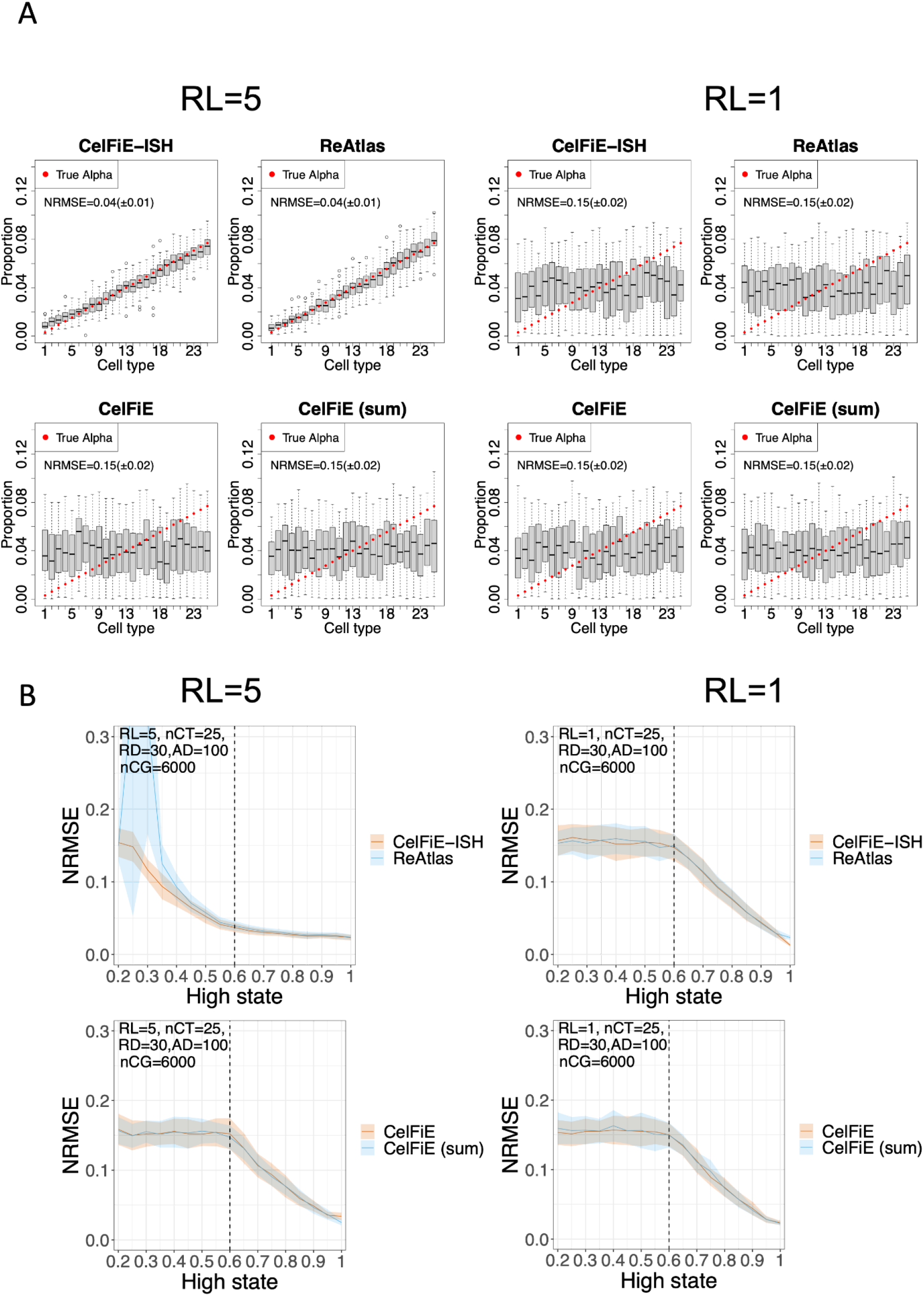
two-state simulations. Simulations using Θ_*low*_=0.1 and Θ_*high*_=0.6, showing the estimated proportion of each cell type (gray box) vs. true proportion (red circle) for each model, using a read length of either RL=5 (left) or RL=1 (right). Each plot is based on 50 replicates simulations, and shows the Normalized Error Normalized error (Normalized Root Mean Square Error or NMRSE) and NRMSE IQR across replicates. (B) NRMSE as a result of varying the Θ_*high*_ state from 0.2 to 1.0 (with [held constant](https://editor.codecogs.com/?latex=%5CTheta_%7Blow%7D#0) at 0.1. Left using read length RL=5, and right using read length RL=1. Shaded area shows standard deviation across 50 replicates.A dotted vertical line shows the condition from panel A above. ReAtlas compared to CelFiE-ISH, CeFiE-Sum compared to CelFiE.

**Figure S3:**
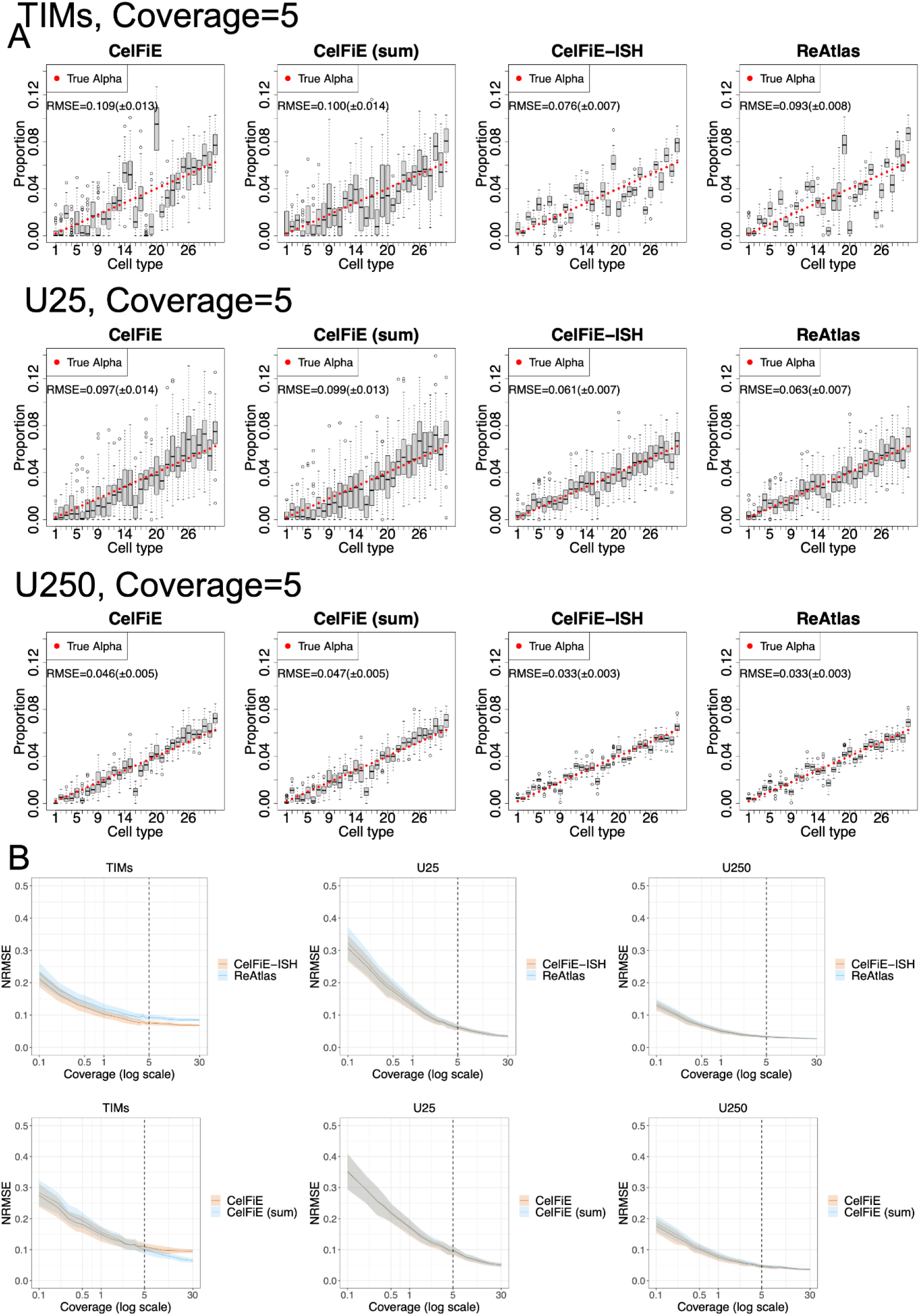
Performance of “Sum” and “Reatlas” model variants in high complexity in silico mixtures. 31 cell types from the Loyfer et al. WGBS methylation atlas were mixed at set proportions to yield a total average read depth (coverage) of 5. (A) Deconvolution results using three different sets of input markers, Caggiano et al. method TIMs (top row), Loyfer et al. U25 (middle row), and Loyfer et al. U250 (bottom row). In each case, estimated proportions are shown as a boxplot of 50 replicates, and true mixture proportions are shown as a red circle. (B) Mixtures were performed for read depths (coverages) from 0.1x to 30x, showing NRMSE for each model and each marker set. Dotted vertical lines indicate the condition from panel A (coverage=5). Shaded areas show standard deviation across 50 replicates.

**Figure S4:**
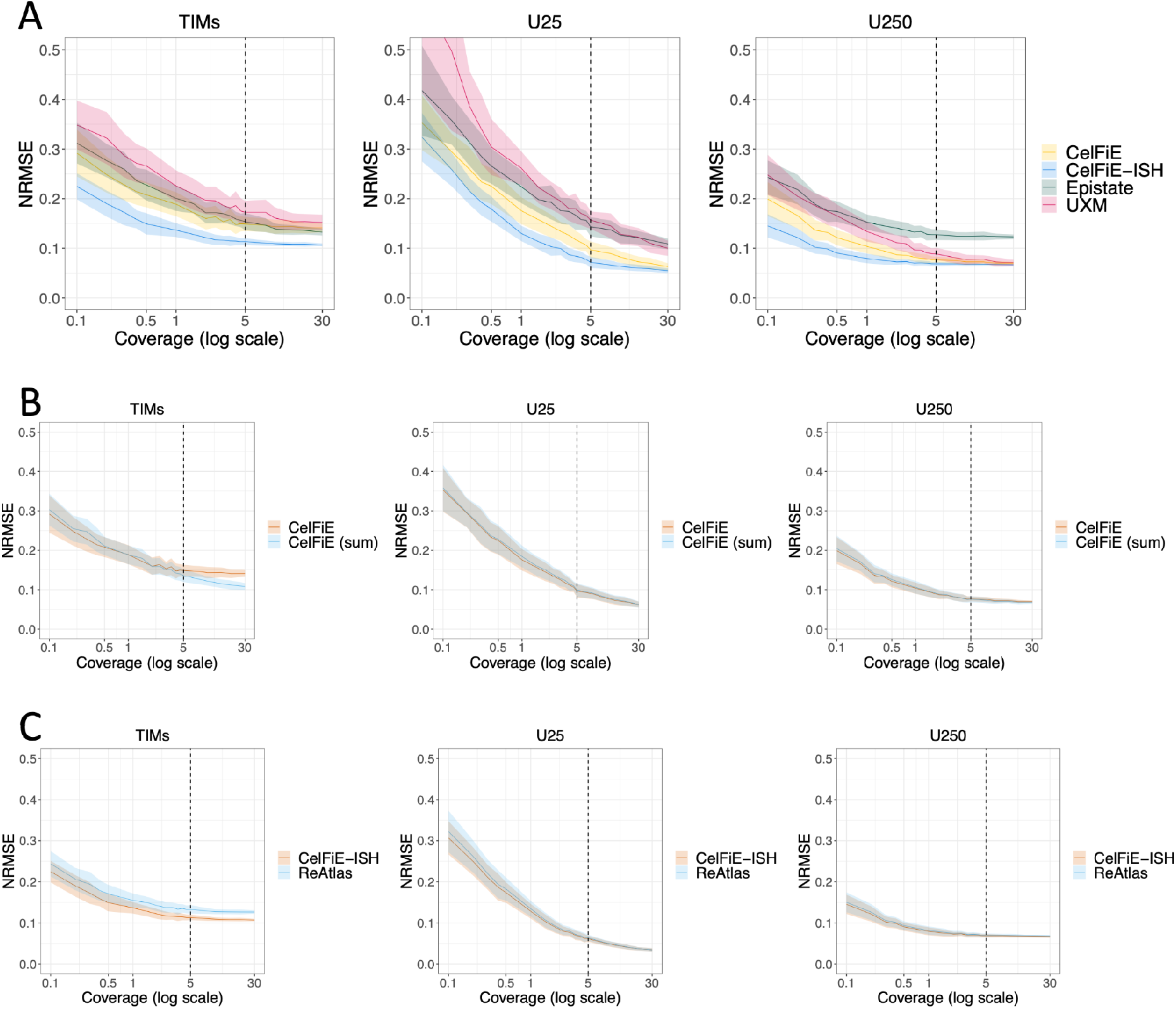
High complexity in silico mixtures with 2nd random cell type shuffling, cell type randomly shuffled. A) NRMSE for each model and each marker set. Dotted vertical lines indicate the condition from figure 3 (coverage=5). B) effect of summing (sum-CelFiE) compared to CelFiE without summing. C) ReAtlas compared to CelFIE-ISH. The cell type order used is: Prostate-Ep, Pancreas-Beta, Lung-Ep-Alveo, Small-Int-Ep, Fallopian-Ep, Eryth-prog, Oligodend, Gastric-Ep, Breast-Basal-Ep, Breast-Luminal-Ep, Liver-Hep, Pancreas-Delta, Bladder-Ep, Blood-NK, Lung-Ep-Bron, Kidney-Ep, Thyroid-Ep, Blood-B, Heart-Cardio, Colon-Ep, Adipocytes, Pancreas-Alpha, Endothelium, Head-Neck-Ep, Blood-Granul, Blood-Mono+Macro, Neuron, Pancreas-Duct, Blood-T, Pancreas-Acinar, Ovary+Endom-Ep.

**Figure S5:**
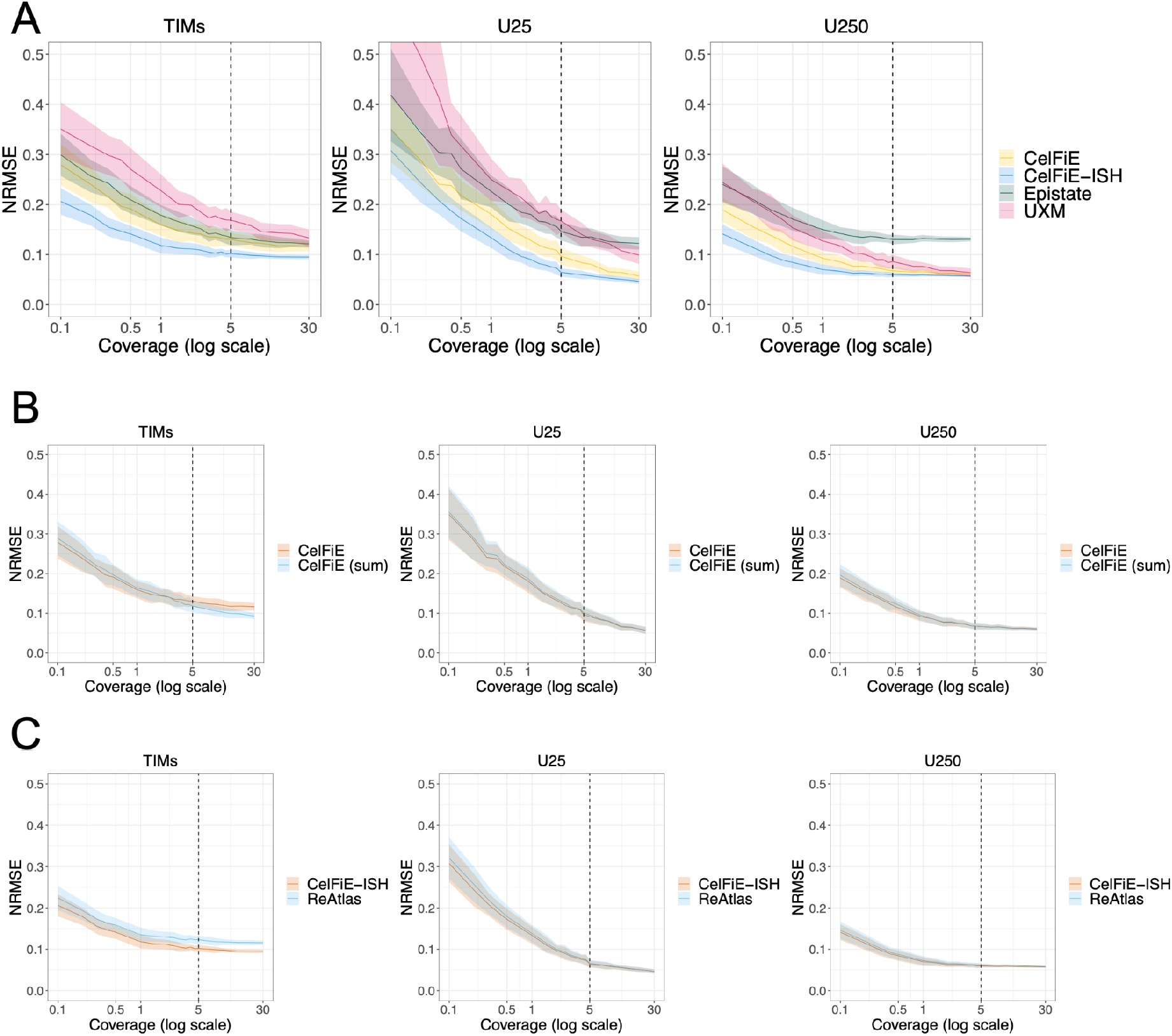
High complexity in silico mixtures with 3rd random cell type shuffling, cell type randomly shuffled. A) NRMSE for each model and each marker set. Dotted vertical lines indicate the condition from figure 3 (coverage=5). B) effect of summing (sum-CelFiE) compared to CelFiE without summing. C) ReAtlas compared to CelFIE-ISH. The cell type order used is: Blood-NK, Head-Neck-Ep, Kidney-Ep, Blood-B, Small-Int-Ep, Breast-Luminal-Ep, Pancreas-Beta, Colon-Ep, Pancreas-Acinar, Gastric-Ep, Prostate-Ep, Breast-Basal-Ep, Eryth-prog, Liver-Hep, Oligodend, Blood-T, Neuron, Fallopian-Ep, Bladder-Ep, Pancreas-Delta, Blood-Granul, Thyroid-Ep, Endothelium, Lung-Ep-Bron, Lung-Ep-Alveo, Ovary+Endom-Ep, Heart-Cardio, Blood-Mono+Macro, Pancreas-Duct, Adipocytes, Pancreas-Alpha.

**Figure S6:**
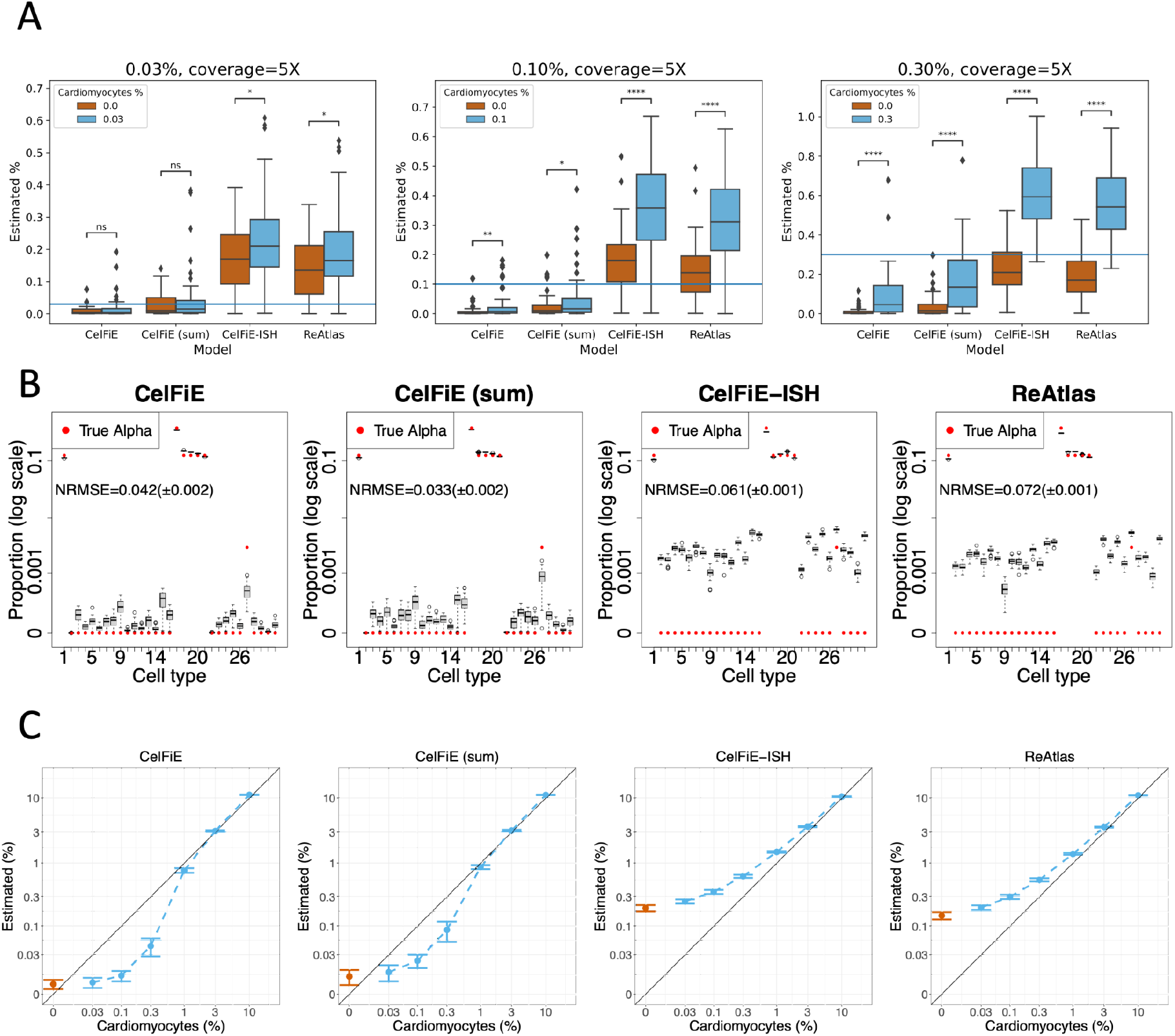
Performance of “Sum” and “Reatlas” model variants in Circulating DNA inspired in silico mixtures. A) Cardiomyocyte samples were added to a background mixture of leukocytes and hepatocytes at varying proportions: 0.03% (left), 0.1% (middle) and 0.3% (right), at a depth of 5X. A one-tailed t-test was performed on the cardiomyocyte estimations of each model against a null mixture with no added cardiomyocytes. *<0.05, **<0.01, ***<0.001, ****<0.00001. B) Full 31-cell type deconvolution of 50 0.3% cardiomyocyte mixtures. In each case, estimated proportions are shown as a boxplot of 50 replicates, and true mixture proportions are shown as a red circle. To display the wide range of true proportions, the Y axis is in log scale.The cell type ordering is listed in the Methods. C) Mixtures of the leukocyte-hepatocyte background were performed for read depth 100,with 0-10% cardiomyocytes. Cardiomyocyte estimates across 50 replicates are shown, with standard deviation. Both axes are in log-scale.

**Figure S7:**
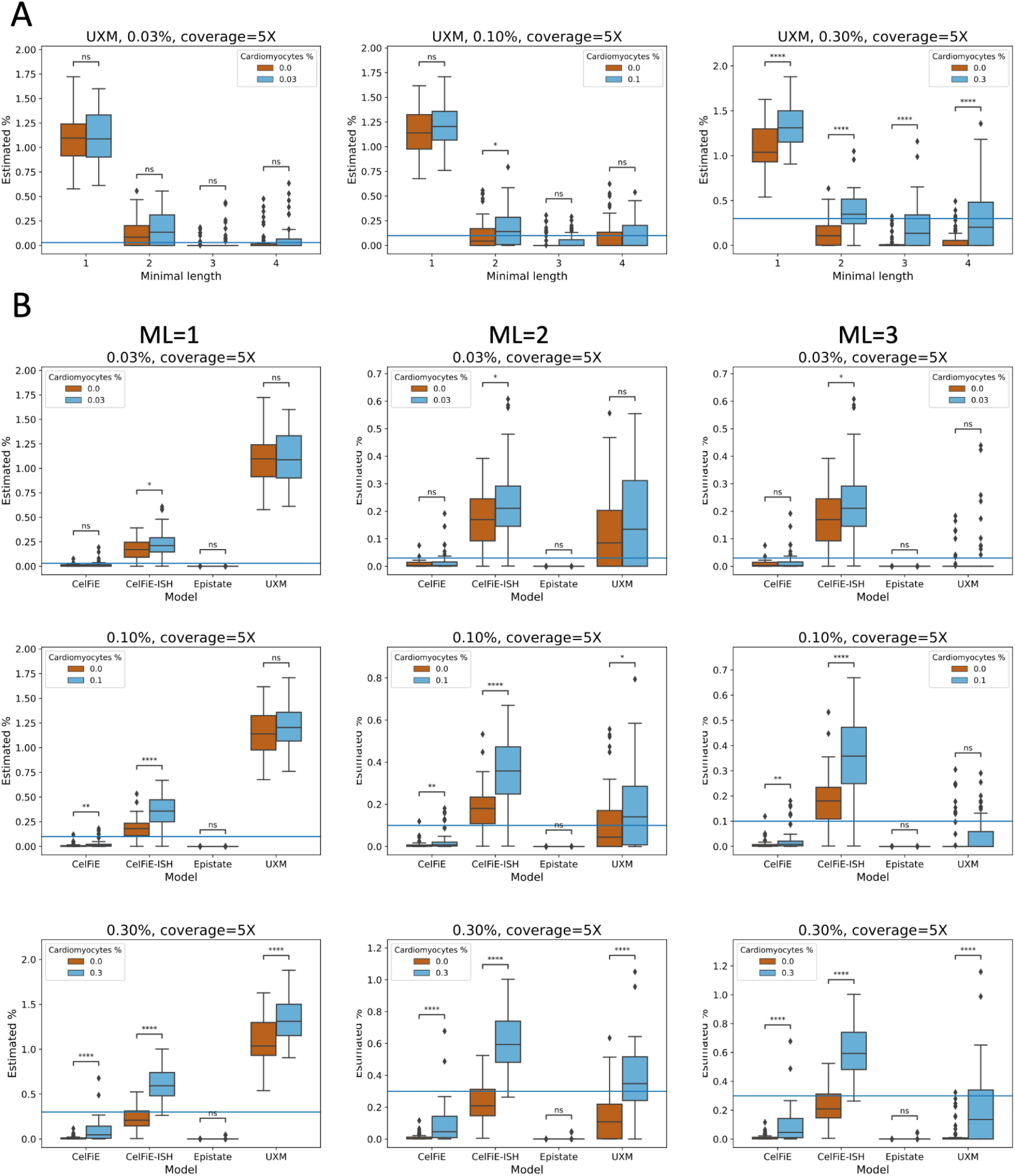
Performance of UXM minimal length adjustment in Circulating DNA inspired in silico mixtures. The settings used for UXM in the Loyfer analyses was a minimum of 4 CpGs per fragment, and a threshold of 25% methylation for U fragments. However, these can be adjusted, and this could have implications on performance, especially in low-coverage settings, where the exclusion of a large proportion of reads could be detrimental to deconvolution. Here, we explore the effect of changing the minimal accepted number of CpGs per fragment (minimal length). The reference atlas remained constant at minimal 4 CpGs. A) UXM with varying thresholds was applied to the same spike-in mixtures shown in Figure 4A. B) UXM minimal length 1 (left), 2 (middle) and 3 (right) along with the other models from Figure 4A, at a target proportion of 0.03% (top), 0.1% (middle) and 0.3% (bottom).

**Figure S8:**
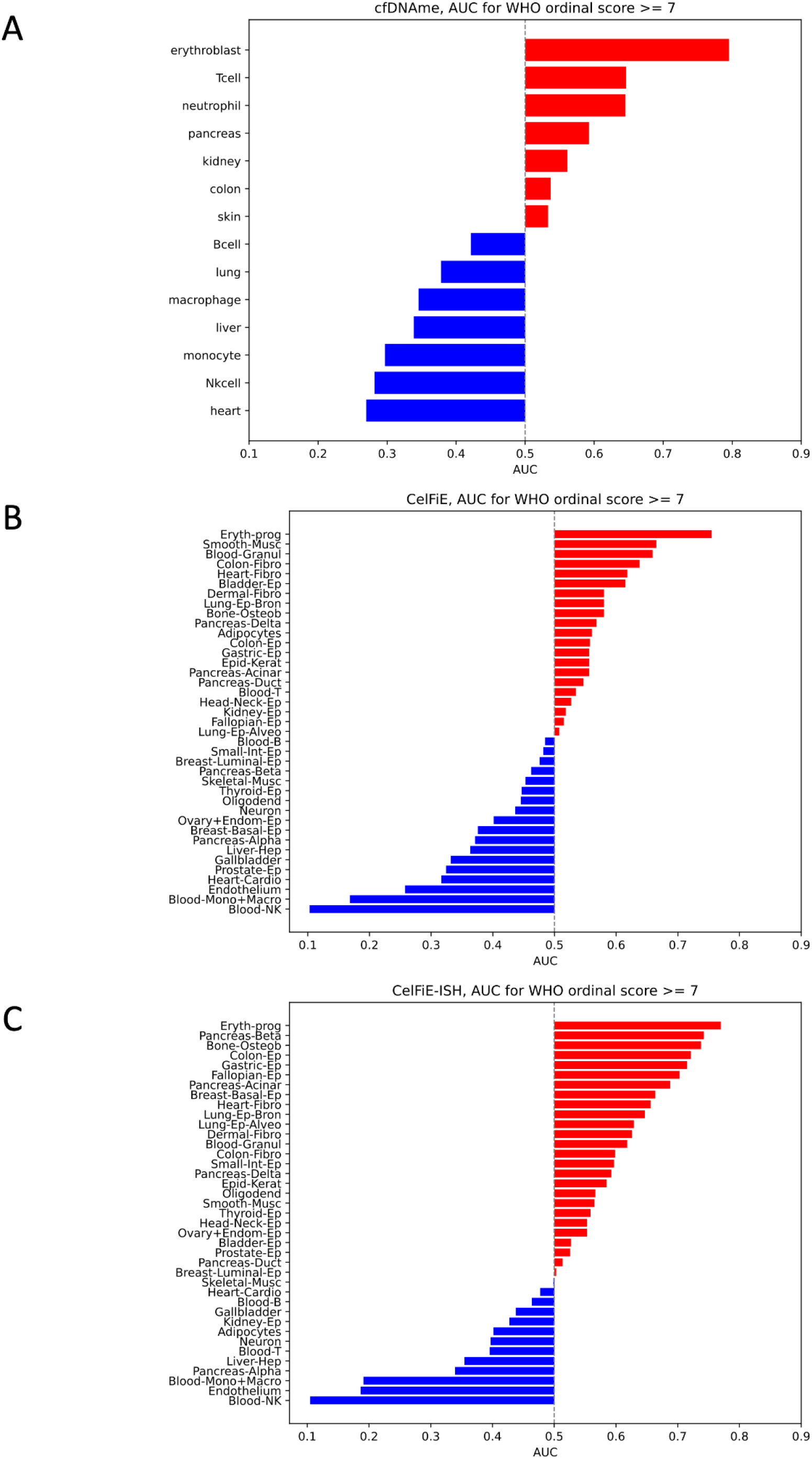
AUC scores for COVID-19 severity. ROC analysis was performed using the proportions for each cell type separately to predict WHO ordinal code >= 7 (patients requiring mechanical ventilation in the ICU). Cell types are ranked by the AUC score. A) cfDNAme, fractions as reported in [28]. AUC of CelFiE (B) and CelFiE-ISH (C) fractions, based on the U250 regions and the 39-cell type Loyfer atlas [1].

**Figure S9:**
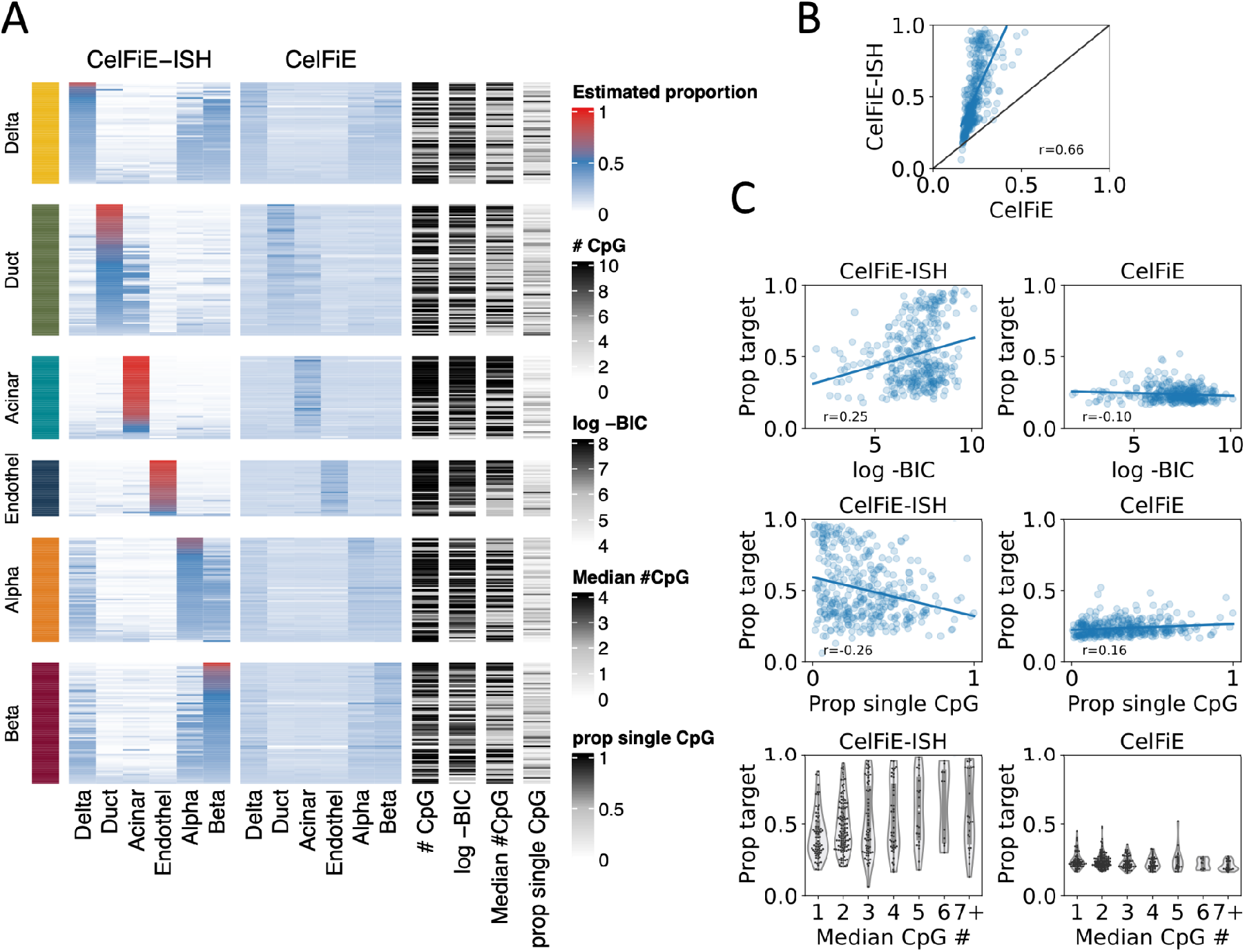
Differential marker contributions to deconvolution accuracy using TIM regions. A) Performance of individual TIM markers in identifying the cell-type specificity between 6 pancreatic cell types. Each group on the horizontal axis shows the accuracy of identifying reads from a specific cell type (i.e. the “yellow” section is performance only on reads derived the pancreatic Delta cell holdout sample). Within each group all TIM markers are displayed as rows, and the percentage of reads identified as each of the 6 cell types in a 6-cell type deconvolution is shown as a heatmap. Rows are ordered by accuracy, i.e. markers that assign the highest fraction of reads to the target cell type are at the top. Additional features of each marker are shown at the right: # CpG: number of CpG sites in the region, log -BIC: log of the Bayesian information criterion, Median # CpG: the median number of CpGs per read, prop single CpG: proportion of reads with a single CpG out of all reads overlapping the region. B) Correlation of the percentage of reads assigned to the target cell type (“Prop target”) between CelFIE and CelFiE-ISH. C) Correlation of the percentage of reads assigned to the target cell type (“Prop target”) between each model and each individual marker features (left column for CelFIE-ISH, right for CelFiE).

**Figure S10:**
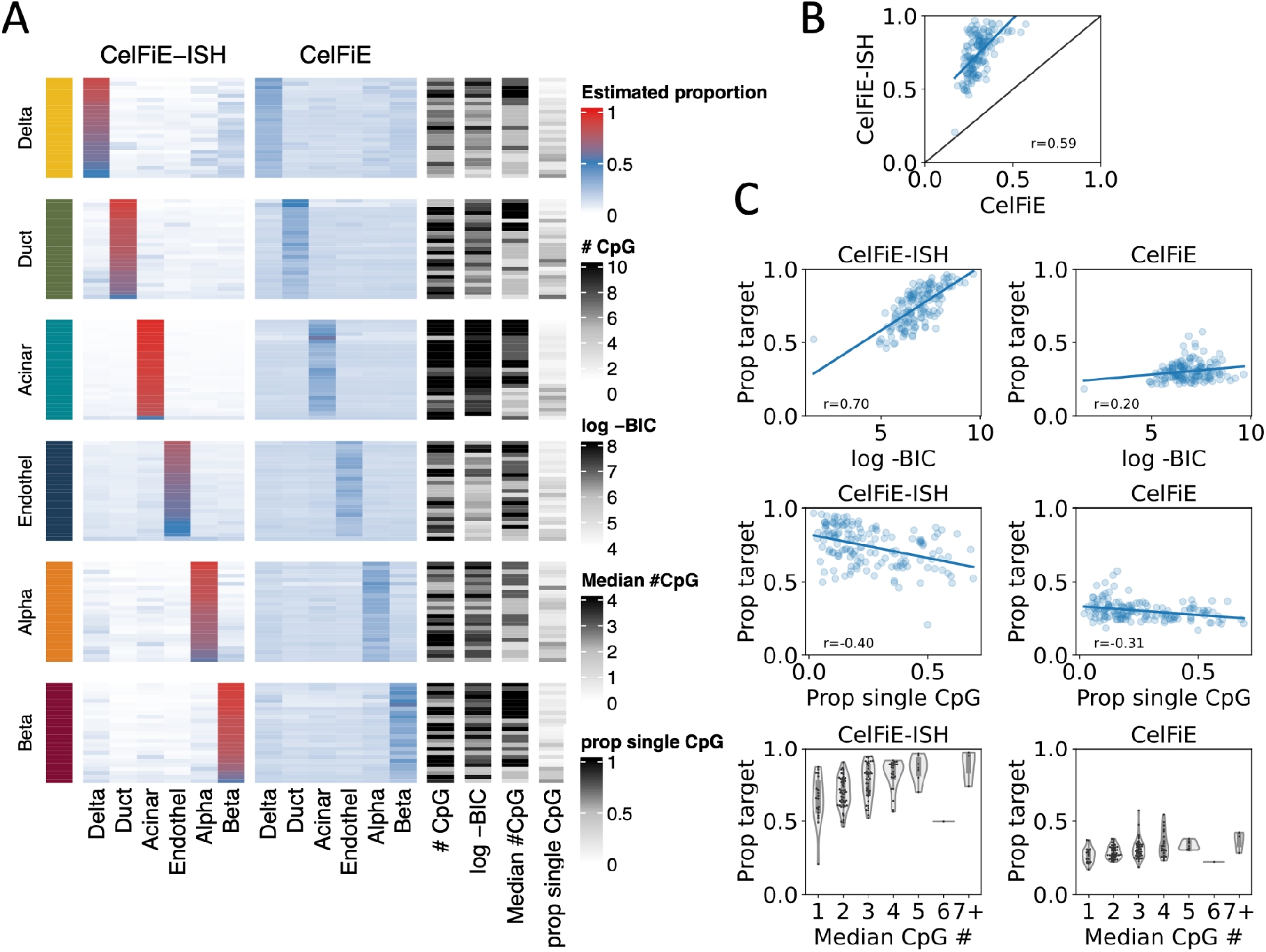
Differential marker contributions to deconvolution accuracy using U25 regions. A) Performance of individual U25 markers in identifying the cell-type specificity between 6 pancreatic cell types. Each group on the horizontal axis shows the accuracy of identifying reads from a specific cell type (i.e. the “yellow” section is performance only on reads derived the pancreatic Delta cell holdout sample). Within each group all U25 markers are displayed as rows, and the percentage of reads identified as each of the 6 cell types in a 6-cell type deconvolution is shown as a heatmap. Rows are ordered by accuracy, i.e. markers that assign the highest fraction of reads to the target cell type are at the top. Additional features of each marker are shown at the right: # CpG: number of CpG sites in the region, log -BIC: log of the Bayesian information criterion, Median # CpG: the median number of CpGs per read, prop single CpG: proportion of reads with a single CpG out of all reads overlapping the region. B) Correlation of the percentage of reads assigned to the target cell type (“Prop target”) between CelFIE and CelFiE-ISH. C) Correlation of the percentage of reads assigned to the target cell type (“Prop target”) between each model and each individual marker features (left column for CelFIE-ISH, right for CelFiE).

### Effect of intra-cell type heterogeneity

In the two-state simulation model, it is possible to alter the association between each cell type and its methylation state, by varying the λ value. For example, a λ of 0.2 for the target cell type would mean that 20% of reads in the target cell type are from Θ_*high*_ and as such are indistinguishable from the background cell types. This is different from varying the beta value parameter: it means there are two subpopulations within the target cell type. This is often seen in WGBS data, and could either indicate contamination from another cell type, or true heterogeneity (both populations belong to the target cell type). To illustrate, say we have an acinar cell sample, covering an acinar marker region with a λ of 20% for acinar cells, and 100% for all other cell types. The methylation states are sufficiently different so that each read can be confidently assigned to a state of origin. With a coverage of 10, 8 reads are from Θ_*low*_ and 2 reads are from Θ_*high*_. The different algorithms treat this hypothetical scenario differently. CelFiE (and Sum-CelFiE) look only at the total methylated and total CpG count and would not differentiate this case from a change in beta value. All the reads would be assigned to acinar cells. CelFiE-ISH (and CelFiE-ISH ReAtlas) would recognise the heterogeneity, and assign the 2 reads to the background cell types. Epistate models this scenario explicitly and would assign the 2 reads to acinar cells. In the case of contamination, where each cell type is actually only associated with one methylation state, CelFiE-ISH would estimate the origins correctly. If however, it is possible to observe both states under one cell type, the Epistate model would estimate the origins correctly.

This effect is heightened with increasing methylation difference and read length, as the states become more distinct. What looks like an increase in error for CelFiE-SH, or a shrinkage effect in the box plots, can also be considered an error of the labels the algorithms are evaluated against.

**Figure S11:**
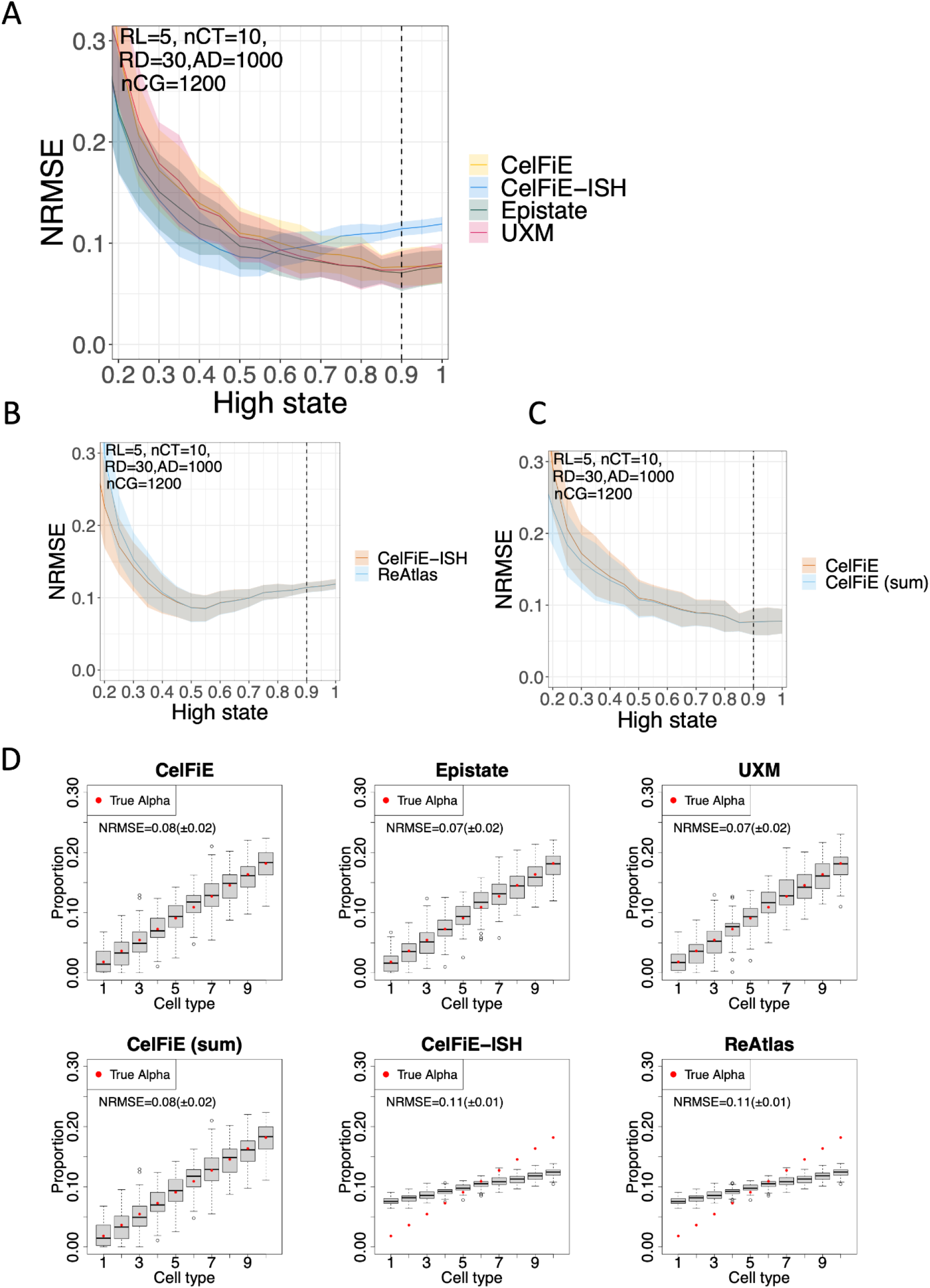
shrinkage effect in simulated conditions. A) NRMSE as a result of varying the Θ_*high*_ state from 0.2 to 1.0, when the is 0.8 (with Θ_*low*_ held constant at 0.1, using read length RL=5). Shaded area shows standard deviation across 50 replicates.A dotted vertical line shows the condition from panel D below. B) ReAtlas compared to CelFiE-ISH. C) CeFiE-Sum compared to CelFiE. D) Shrinkage at Θ_*high*_ =0.9.

