## Supplementary Note for "Multi-cell type deconvolution using a probabilistic model for single-molecule DNA methylation haplotypes"

### The CelFiE-ISH Model

#### 1 Reference Atlas

The reference atlas consists of one matrix $\beta_{t,m}$, with the probability of methylation for cell type $t$ at position $m$. In this model we do not re-estimate the atlas at each iteration.

#### 2 Mixture

The mixture is one matrix $X$, with dimensions $C$ reads over $M$ CpG sites.

#### 3 Likelihood

The observed data likelihood is:

$$P(x|\alpha,\beta) = \prod_c \sum_t \alpha_t P(x_c|\beta_t) = \prod_c \sum_t \alpha_t \prod_m \beta_{t,m}^{x_{c,m}} (1-\beta_{t,m})^{1-x_{c,m}} \tag{1}$$

The observed data log-likelihood is:

$$logP(x|\alpha,\beta) = \sum_c log(\sum_t \alpha_t \prod_m \beta_{t,m}^{x_{c,m}} (1-\beta_{t,m})^{1-x_{c,m}}) = \sum_c logsumexp\left\{ log(\alpha_t \prod_m \beta_{t,m}^{x_{c,m}} (1-\beta_{t,m})^{1-x_{c,m}}) \right\} = \sum_c logsumexp\left\{ log(\alpha_t) + \sum_m x_{c,m} log(\beta_{t,m}) + (1-x_{c,m}) log(1-\beta_{t,m}) \right\} \tag{2}$$

The complete data likelihood is:

$$P(x,z|\alpha,\beta) = P(x|z,\beta)P(z|\alpha)$$

(3)

Where the first term is

$$log(P(x|z,\beta)) = \sum_{t,c,m} log\left[\beta_{t,m}^{z_{t,c}x_{c,m}}(1-\beta_{t,m})^{z_{t,c}(1-x_{c,m})}\right[\right]$$
$$= \sum_{t,c,m} z_{t,c}[x_{c,m}log(\beta_{t,m}) + (1-x_{c,m})log(1-\beta_{t,m})]$$

(4)

and the second term is

$$log(P(z|\alpha)) = \sum_{t,c} log(\alpha_t^{z_{t,c}}) = \sum_{t,c} z_{t,c}log(\alpha_t)$$

(5)

### 4 Q function

As $z$ in unknown, we define $\tilde{p}$ as the probability of $z$:

$$P(z_{t,c} = 1|\alpha,\beta) =: \tilde{p}_{t,c}$$

Q is the expected value of the log-likelihood function.

At iteration $i$, the Q-function is:

$$Q_i = \mathbb{E}_{z|x,\alpha^i,\beta}(logP(x,z|\alpha,\beta)) =$$
$$\sum_{t,c}\tilde{p}_{t,c}^i \sum_m [x_{c,m}log(\beta_{t,m}) + (1-x_{c,m})log(1-\beta_{t,m})] +$$
$$\sum_{t,c}\tilde{p}_{t,c}^i log(\alpha_t)$$

(6)

#### 5 E-step

In the E-step we estimate the latent variable $z$ and use it to define the Q function.

$$P(z_{t,c} = 1 | x_c, \beta, \alpha) = \frac{\alpha_t \prod_m \beta_{t,m}^{x_{m,c}} (1-\beta_{t,m})^{1-x_{m,c}}}{\sum_k \alpha_k \prod_m \beta_{k,m}^{x_{m,c}} (1-\beta_{k,m})^{1-x_{m,c}}} =: \tilde{p}_{t,c} \tag{7}$$

#### 6 M-step

In the M-step we maximize the Q function, holding the estimate for the latent variable $z$ constant and maximizing $\alpha$.

$$\alpha_t = \frac{\sum_c \tilde{p}_{t,c}}{C}$$

### The CelFiE-ISH ReAtlas Model

#### 7 Reference Atlas

The reference atlas consists of two matrices, $Y_{t,m}$ and $D^Y_{t,m}$, with the number of methylated and total reads for cell type $t$ at position $m$ respectively. We assume $Y_{t,m}$ is drawn from a Binomial distribution with $\beta_{t,m}$ being the true methylation probability and $D^Y_{t,m}$ being the number of trials. We re-estimate the atlas at each iteration.

#### 8 Mixture

The mixture is one matrix $X$, with dimensions $C$ reads over $M$ CpG sites.

#### 9 Likelihood

The observed data likelihood is:

$$P(x|\alpha,\beta) = P(x|\alpha,\beta)P(Y|\beta) = \prod_c \sum_t \alpha_t P(x_c|\beta_t) P(Y|\beta) =$$

$$\prod_c \left\{ \sum_t \alpha_t \prod_m \beta_{t,m}^{x_{c,m}} (1-\beta_{t,m})^{1-x_{c,m}} \right\} \prod_t \prod_m \left\{ \beta_{t,m}^{Y_{t,m}} (1-\beta_{t,m})^{D^{Y_{t,m}} - Y_{t,m}} \right\} \tag{8}$$

The observed data log-likelihood is:

$$logP(x|\alpha,\beta) = \sum_c log(\sum_t \alpha_t \prod_m \beta_{t,m}^{x_{c,m}}(1-\beta_{t,m})^{1-x_{c,m}}) + log(P(Y|\beta)) =$$

$$\sum_c logsumexp\left\{log(\alpha_t \prod_m \beta_{t,m}^{x_{c,m}}(1-\beta_{t,m})^{1-x_{c,m}})\right\} + log(P(Y|\beta)) =$$

$$\sum_c logsumexp\left\{log(\alpha_t) + \sum_m x_{c,m}log(\beta_{t,m}) + (1-x_{c,m})log(1-\beta_{t,m})\right\} + log(P(Y|\beta)) =$$

$$\sum_c logsumexp\left\{log(\alpha_t) + \sum_m x_{c,m}log(\beta_{t,m}) + (1-x_{c,m})log(1-\beta_{t,m})\right\} +$$

$$\sum_{t,m}\left\{Y_{t,m}log\beta_{t,m} + (D^{Y_{t,m}} - Y_{t,m})log(1-\beta_{t,m})\right\}$$

(9)

The complete data likelihood is:

$$P(x,z,Y|\alpha,\beta) = P(x|z,\beta)P(z|\alpha)P(Y|\beta)$$

(10)

The first term is

$$log(P(x|z,\beta)) = \sum_{t,c,m} log\left[\beta_{t,m}^{z_{t,c}x_{c,m}}(1-\beta_{t,m})^{z_{t,c}(1-x_{c,m})}[\right]$$

$$= \sum_{t,c,m} z_{t,c}[x_{c,m}log(\beta_{t,m}) + (1-x_{c,m})log(1-\beta_{t,m})]$$

(11)

The second term is

$$log(P(z|\alpha)) = \sum_{t,c} log(\alpha_t^{z_{t,c}}) = \sum_{t,c} z_{t,c}log(\alpha_t)$$

(12)

The third term is

$$log(P(Y|\beta)) = \sum_{t,m} Y_{t,m} log\beta_{t,m} + (D^{Y_{t,m}} - Y_{t,m}) log(1 - \beta_{t,m})$$

(13)

#### 10 Q function

As $z$ in unknown, we define $\tilde{p}$ as the probability of $z$:

$$P(z_{t,c} = 1|\alpha, \beta) =: \tilde{p}_{t,c}$$

Q is the expected value of the log-likelihood function.

At iteration $i$, the Q-function is:

$$Q_i = \mathbb{E}_{z|x,\alpha^i,\beta^i}(logP(x, z, Y|\alpha, \beta)) =$$
$$\sum_{t,c} \tilde{p}^i_{t,c} \sum_m [x_{c,m} log(\beta_{t,m}) + (1 - x_{c,m}) log(1 - \beta_{t,m})] +$$
$$\sum_{t,c} \tilde{p}^i_{t,c} log(\alpha_t) +$$
$$\sum_{t,m} Y_{t,m} log\beta_{t,m} + (D^{Y_{t,m}} - Y_{t,m}) log(1 - \beta_{t,m})$$

(14)

#### 11 E-step

In the E-step we estimate the latent variable $z$ and use it to define the Q function.

$$P(z_{t,c} = 1|x_c, \beta, \alpha) = \frac{\alpha_t \prod_m \beta_{t,m}^{x_{m,c}} (1 - \beta_{t,m})^{1-x_{m,c}}}{\sum_k \alpha_k \prod_m \beta_{k,m}^{x_{m,c}} (1 - \beta_{k,m})^{1-x_{m,c}}} =: \tilde{p}_{t,c}$$

(15)

#### 12 M-step

In the M-step we maximize the Q function, holding the estimate for the latent variable $z$ constant and maximizing $\alpha$.

$$\alpha_t = \frac{\sum_c \tilde{p}_{t,c}}{C}$$

Next, we re-estimate the atlas:

$$\beta_{t,m} = \frac{Y_{t,m}+\sum_c \tilde{p}_{t,c}x_{c,m}}{D^{Y_{t,m}}+\sum_c \tilde{p}_{t,c}}$$

(16)

### The Epistate Model

At every marker region, reads are drawn from one of two possible epistates: $\theta_{high}$ and $\theta_{low}$. Each epistate consists of a set of binomial distributions $\theta = \{\theta_1, \theta_2, ..., \theta_m\}$, one per CpG site covered by the marker region. $\theta_{high}$ is arbitrarily defined to be the epistate with higher mean methylation. Cell types differ by the probability of observing each epistate in each region.

### 13 Reference Atlas

The reference atlas consists of one matrix $\lambda_{t,c}$, with the probability of observing $\theta_{high}$ under cell type $t$ at read $c$. Within a genomic region $\lambda$ does not vary between reads, leaving $\lambda_t$. Additionally, for every position we know $\theta_{high,m}$ and $\theta_{low,m}$ (see below). The overall probability of methylation per position is:

$$\beta_{t,m} = \lambda_t\theta_{high,m} + (1-\lambda_t)\theta_{low,m}$$

### 14 Mixture

The mixture is one matrix $X$, with dimensions $C$ reads over $M$ CpG sites.

### 15 Likelihood

The observed data likelihood is:

$$P(x|\alpha, \theta_{high}, \theta_{low}, \lambda) = \prod_c \sum_t \alpha_t \left\{ \lambda_{t,c} \prod_m \left[ \theta_{high}^{x_{c,m}} (1-\theta_{high})^{1-x_{c,m}} \right] + (1-\lambda_{t,c}) \prod_m \left[ \theta_{low}^{x_{c,m}} (1-\theta_{low})^{1-x_{c,m}} \right] \right\}$$

(17)

The observed data log-likelihood is:

$$logP(x|\alpha, \theta_{high}, \theta_{low}, \lambda) = \sum_c log(\sum_t \alpha_t \Bigg\{ \lambda_{t,c} \prod_m \Bigg[ \theta_{high}^{x_{c,m}} (1-\theta_{high})^{1-x_{c,m}} \Bigg] +$$
$$(1-\lambda_{t,c}) \prod_m \Bigg[ \theta_{low}^{x_{c,m}} (1-\theta_{low})^{1-x_{c,m}} \Bigg] \Bigg\}) =$$
$$\sum_c logsumexp_t \Bigg\{ log(\alpha_t) + log(\lambda_{t,c} \prod_m \Bigg[ \theta_{high}^{x_{c,m}} (1-\theta_{high})^{1-x_{c,m}} \Bigg] +$$
$$(1-\lambda_{t,c}) \prod_m \Bigg[ \theta_{low}^{x_{c,m}} (1-\theta_{low})^{1-x_{c,m}} \Bigg]) \Bigg\} =$$
$$\sum_c logsumexp_t \Bigg\{ log(\alpha_t) + logsumexp \Bigg\{ log(\lambda_{t,c}) + \sum_m \Bigg[ x_{c,m} log(\theta_{high}) + (1-x_{c,m}) log(1-\theta_{high}) \Bigg],$$
$$log(1-\lambda_{t,c}) + \sum_m \Bigg[ x_{c,m} log(\theta_{low}) + (1-x_{c,m}) log(1-\theta_{low}) \Bigg] \Bigg\} \Bigg\}$$

(18)

$z$ is the indicator for $\alpha$ and $\mu$ is the indicator for $\lambda$. The complete data likelihood is:

$$P(x, z, \mu|\alpha, \theta_{high}, \theta_{low}, \lambda) = P(x|\mu, \theta_{high}, \theta_{low}) P(z|\alpha) P(\mu|z, \lambda)$$

(19)

The first term is

$$log(P(x|\mu, \theta_{high}, \theta_{low})) = log(\prod_c \prod_m \Bigg[ \theta_{high,m}^{\mu_c x_{c,m}} (1-\theta_{high,m})^{\mu_c(1-x_{c,m})}$$
$$\theta_{low,m}^{(1-\mu_c)x_{c,m}} (1-\theta_{low,m})^{(1-\mu_c)(1-x_{c,m})} \Bigg]) =$$
$$\sum_{c,m} \Bigg[ \mu_c x_{c,m} log(\theta_{high,m}) + \mu_c (1-x_{c,m}) log(1-\theta_{high,m}) +$$
$$(1-\mu_c) x_{c,m} log(\theta_{low,m}) + (1-\mu_c)(1-x_{c,m}) log(1-\theta_{low,m}) \Bigg]$$

(20)

The second term is

$$log(P(z|\alpha)) = \sum_{t,c} log(\alpha_t^{z_{t,c}}) = \sum_{t,c} z_{t,c} log(\alpha_t) \tag{21}$$

The third term is

$$log(P(\mu|z,\lambda)) = log(\prod_t \prod_c \lambda_{t,c}^{z_{t,c}\mu_c}(1-\lambda_{t,c})^{z_{t,c}(1-\mu_c)}) = \sum_{t,c}\Big[z_{t,c}\mu_c log(\lambda_{t,c}) + z_{t,c}(1-\mu_c) log(1-\lambda_{t,c})\Big] \tag{22}$$

#### 16 Q function

As $z$ in unknown, we define $\tilde{p}$ as the posterior probability of $z$:

$$P(z_{t,c} = 1|\alpha, x) =: \tilde{p}_{t,c}$$

Similarly,

$$P(\mu_c = 1|z, x) =: \tilde{q}_c$$

Note that $\lambda$, $\theta_{high}$, $\theta_{low}$ and by extension $\beta$ are always given and not re-estimated. For simplicity, we left them out of the conditional statements.
Q is the expected value of the log-likelihood function.

At iteration $i$, the Q-function is:

$$Q_i = \mathbb{E}_{z,\mu|x,\alpha^i,\lambda,\theta_{high},\theta_{low}}(logP(x,z,\mu|\alpha^i,\theta_{high},\theta_{low},\lambda)) = \sum_{t,c}\Big\{\tilde{p}_{t,c}\tilde{q}_c \sum_m \Big[x_{c,m} log(\theta_{high,m}) + (1-x_{c,m}) log(1 [illegible]$$

(23)

#### 17 E-step

In the E-step we estimate the latent variables $z$ and $\mu$ and use them to define the Q function.

$$P(\mu_c = 1|x,\alpha) = \sum_t P(z_{t,c} = 1|x,\alpha_t) P(\mu_c = 1|z_{t,c} = 1, x, \alpha) = \sum_t \tilde{p}_{t,c} P(\mu_c = 1|z_{t,c} = 1, x) \propto \sum_t \tilde{p}_{t,c} P(x|\mu_c = 1, z_{t,c} = 1) P(\mu_c = 1|z_{t,c} = 1) = \sum_t \tilde{p}_{t,c} P(x|\mu_c = 1) P(\mu_c = 1|z_{t,c} = 1) = \sum_t \tilde{p}_{t,c}\lambda_t P(x|\mu_c = 1) = \sum_t \tilde{p}_{t,c}\lambda_t \prod_m \theta_{high}^{x_{c,m}}(1-\theta_{high})^{1-x_{c,m}} \tag{24}$$

Since $\mu$ can only take on two values, we constrain

$$P(\mu_c = 1|x, \tilde{p}) + P(\mu_c = 0|x, \tilde{p}) = 1$$

As above:

$$P(\mu_c = 0|x, \tilde{p}) = \sum_t \tilde{p}_{t,c}(1-\lambda_t) \prod_m \theta_{low}^{x_{c,m}} (1-\theta_{low})^{1-x_{c,m}}$$

Finally:

$$P(\mu_c = 1|x, \alpha) = \frac{\sum_t \tilde{p}_{t,c}\lambda_t \prod_m \theta_{high}^{x_{c,m}}(1-\theta_{high})^{1-x_{c,m}}}{\sum_t \tilde{p}_{t,c}\lambda_t \prod_m \theta_{high}^{x_{c,m}}(1-\theta_{high})^{1-x_{c,m}} + \sum_t \tilde{p}_{t,c}(1-\lambda_t)\prod_m \theta_{low}^{x_{c,m}}(1-\theta_{low})^{1-x_{c,m}}} \tag{25}$$

We do the same for $z$:

$$\begin{aligned} P(z_{t,c} = 1|x, \alpha_t) \propto P(x|z_{t,c} = 1, \alpha_t)P(z_{t,c} = 1|\alpha_t) &= \Big[\lambda_{t,c}P(x|\mu_c = 1) + (1-\lambda_{t,c})P(x|\mu_c = 0)\Big]\alpha_t \\ &= \alpha_t\lambda_{t,c}\prod_m \Big[\theta_{high}^{x_{c,m}}(1-\theta_{high})^{1-x_{c,m}}\Big] + \alpha_t(1-\lambda_{t,c})\prod_m \Big[\theta_{low}^{x_{c,m}}(1-\theta_{low})^{1-x_{c,m}}\Big] \end{aligned} \tag{26}$$

Then normalize so that every read comes from a cell type.

#### 18 M-step

In the M-step we maximize the Q function, holding the estimate for the latent variables constant and maximizing $\alpha$. The only term in the Q function with $\alpha$ is identical to CelFiE and CelFiE+, so the maximization step is the same.

$$\alpha_t = \frac{\sum_c \tilde{p}_{t,c}}{C}$$

### Estimating Epistates in the Reference Atlas

For each marker region in the Epistate reference, we estimate $\Theta_{high}$, $\Theta_{low}$ and $\lambda_t$. First, we jointly examine all reads from the entire reference dataset. We assume each read is associated with either $\Theta_{high}$ or $\Theta_{low}$. $\upsilon_j$ is the prior probability for epistate $j \in [1, 2]$. At the expectation step, we update the posterior probability of each read $P_{j,c}$ given $\Theta$. At the maximization step, we estimate the hidden state $\Theta$, and $\upsilon_j$.

#### 19 Likelihood

The observed data likelihood is:

$$P(x|\Theta_{high}, \Theta_{low}, \upsilon) = \prod_c \sum_{j=1}^{2} \upsilon_j \Big[\prod_m \theta_j^{x_{c,m}}(1-\theta_j)^{1-x_{c,m}}\Big]$$

#### Expectation

$$P_{j,c} = \frac{\upsilon_j \prod_m \theta_{m,j}^{x_{c,m}} (1-\theta_{k,j})^{1-x_{c,m}}}{\sum_{j=1}^{2} \upsilon_j \prod_m \theta_{m,j}^{x_{c,m}} (1-\theta_{k,j})^{1-x_{c,m}}}$$

#### Maximization

$$\theta_{m1} = \frac{pseudocount + \sum_c P_{1,c} x_{c,m}}{2 * pseudocount + \sum_c P_{1,c}}$$

$$\upsilon_1 = \frac{pseudocount + \sum_c P_{1,c}}{2 * pseudocount + C}$$

Then, we split the reference by cell type. For each cell type, $\lambda$ if the probability of observing $\Theta_{high}$. For each subset:

$$\lambda_t = \frac{\sum_c P_{1,c}}{C}$$

#### WGBS Data Processing

In order to convert BAM files to the Biscuit epiread format, we first generated a SNP file from the VCF files requiring $GQ \geq 15$ for positions overlapping a dbSNP common allele, and requiring $GQ \geq 60$ for all other positions. DbSNP common allele table was downloaded from UCSC for the hg19 assembly, and was processed with:
https://github.com/ekushele/methylseq/blob/master/bin/processUcscDbsnp.pl.

From the processed file, we included only 'snv' records. The formatted-snv file was zipped and indexed with the `tabix -s 1 -b 2 -e 3` command. This file was passed to bcftools annotate (v1.9) to annotate the header of VCF files:
`bcftools annotate WHITELIST -O z -a {COMMON_DBSNP_FILE} -h common_dbsnp.hdr -c CHROM,FROM,TO,TYPE,COMMON_SOME,COMMON_ALL,REF_MIN,ALT_MIN,REF_DBSNP, ALT_DBSNP,REF_ALL,ALT_ALL,RSID,MAX_MAF {VCF_FILE}`.
(common_dbsnp.hdr can be found at:
https://github.com/ekushele/methylseq/blob/master/assets/common_dbsnp.hdr).

The redhead file was indexed with `tabix -p vcf`. From the re-headed files, we included variants with $GQ \geq 60$ for heterozygous variants for positions not overlapping the COMMON_DBSNP_FILE with `bcftools view -O z -i 'ALT!="N" & ALT!="." & ((COUNT(GT=="0/1")` $\geq 1\&COMMON\_ALL == 1\&MAX\_MAF \geq 0.05) \,|\, (COUNT(GT == "0/1"\&GQ \geq 60) \geq 1))'\{REHEAD\_VCF\} > \{DBSNP\_HET60\}$.

{DBSNP_HET60} was indexed with `tabix -p vcf.` For all other variants, we excluded variants below 10 and parsed the file to be in bed format with the

following command:
`bcftools query -u -i 'GT="0/1" & GQ` $\geq 10' - -format'$
$\%CHROM\%POS\%POS\%REF\%ALT[\%GT\%GQ\%SP\%AC\%AF1]\%RSID\%$
$COMMON\_ALL\%MAX\_MAF\%REF\_MIN\%ALT\_MIN'\{DBSNP\_HET60\}|$
$awk - vOFS = "nt"'\{\$2 = \$2 - 1; print\}' > \{SNP\_FILE\}$.
Then, blacklist regions were excluded from BAM files with the command `bedtools intersect`($v2.29.1$) using the BAM and a whitelist as input files, and additional command line arguments`'-ubam -f 1.0'`.

Epiread files were produced with the `biscuit epiread` command for whitelist-BAM files where a SNP file was given as input to the `-B` argument: `'-B SNP_FILE'`. The epiread files were sorted by names using the command `'-k2,2 -k1,1 -k4,4 -k3,3n'` , and they were converted to a bed-like format, merging paired-end epiread records together using the script available at
`https://github.com/ekushele/methylseq/blob/master/bin/epiread_pairedEnd_convertion`
in debug mode.

The CpG file was downloaded from the Biscuit QC assets release page:
`https://github.com/huishenlab/biscuit/releases`
These merged files were sorted by position using the command `sort -k1,1Vf -k 2,2n -k 3,3n` and then tabixed using the `'tabix -0 -p bed'` command. The original epireads (before merging) were sorted with `sort -k1,1Vf -k5,5V` and tabixed with `tabix -0 -s 1 -b 5 -e 5`.
